# Swedish Nerve Growth Factor Mutation (NGF^R100W^) Defines a Role for TrkA and p75NTR in Nociception

**DOI:** 10.1101/266643

**Authors:** Kijung Sung, Luiz F. Ferrari, Wanlin Yang, ChiHye Chung, Xiaobei Zhao, Yingli Gu, Suzhen Lin, Kai Zhang, Bianxiao Cui, Matthew L. Pearn, Michael T. Maloney, William C. Mobley, Jon D. Levine, Chengbiao Wu

## Abstract

Nerve growth factor (NGF) exerts multiple functions on target neurons throughout development. The recent discovery of a point mutation leading to a change from arginine to tryptophan at residue 100 in the mature NGFβ sequence (NGF^R100W^) in patients with hereditary sensory and autonomic neuropathy, type V (HSAN V), made it possible to distinguish the signaling mechanisms that lead to two functionally different outcomes of NGF: trophic versus nociceptive. We performed extensive biochemical, cellular and live imaging experiments to examine the binding and signaling properties of NGF^R100W^. Our results show that, similar to the wildtype NGF (wtNGF), the naturally occurring NGF^R100W^ mutant was capable of binding to and activating the TrkA receptor and its downstream signaling pathways to support neuronal survival and differentiation. However, NGF^R100W^ failed to bind and stimulate the 75kD neurotrophic factor receptor (p75^NTR^)-mediated signaling cascades (i.e. the RhoA-Cofilin pathway). Intraplantar injection of NGF^R100W^ into adult rats induced neither TrkA-mediated thermal nor mechanical acute hyperalgesia, but retained the ability to induce chronic hyperalgesia based on agonism for TrkA signaling. Taken together, our studies provide evidence that NGF^R100W^ retains trophic support capability through TrkA and one aspect of its nociceptive signaling, but fails to engage p75^NTR^ signaling pathways. Our findings suggest that wtNGF acts through TrkA to regulate the delayed priming of nociceptive responses. The integration of both TrkA and p75^NTR^ signaling thus appears to regulate neuroplastic effects of NGF in peripheral nociception.

**Significance Statement:** In the present study, we characterized the naturally occurring NGF^R100W^ mutant that is associated with hereditary sensory and autonomic neuropathy, type V. We have demonstrated for the first time that NGF^R100W^ retains trophic support capability through TrkA but fails to engage p75^NTR^ signaling pathways. Furthermore, following Intraplantar injection into adult rats, NGF^R100W^ induced neither thermal nor mechanical acute hyperalgesia, but retained the ability to induce chronic hyperalgesia. We have also provided evidence that the integration of both TrkA-and p75^NTR^-mediated signaling thus appears to regulate neuroplastic effects of NGF in peripheral nociception. Our study with NGF^R100W^ suggests that it is possible to uncouple trophic effect from nociceptive function, both induced by wildtype NGF.

**Abbreviations:**

NGF: nerve growth factor;
NGF^R100W^: NGF mutation with a change from tryptophan (W) to arginine (R) at the 100 residue.
TrkA: Tropomyosin receptor kinase A;
p75^NTR^: the 75kD neurotrophic factor receptor;
HSAN V: hereditary sensory and autonomic neuropathy, type V;
BFCN: basal forebrain cholinergic neurons;
PNS: peripheral nervous system;
CNS: central nervous system;
BDNF: brain-derived neurotrophic factor;
NT: neurotrophin;
TrkB: Tropomyosin receptor kinase B;
TrkC: Tropomyosin receptor kinase C;
RhoA: Ras homolog gene family, member A;
NF-κB: nuclear factor kappa B;
Akt: Protein kinase B;
JNK: c-Jun N-terminal kinases;
AD: Alzheimer’s disease;
CSF: cerebrospinal fluid;
HIV: human immunodeficiency virus
CIPA: congenital insensitivity to pain with anhidrosis;
ERK: extracellular signal-related kinase;
PI3K: phosphatidylinositol 3-kinase;
PLCγ: and phospholipase Cγ;
PGE_2_: Prostaglandin E2;
PCR: polymerase chain reaction;
GFP: green fluorescent protein;
QD: quantum dots;
HEK293FT: Human embryonic kidney 293FT cell line;
KKE: NGF mutant protein with mutation at: <u>K</u>32A/<u>K</u>34A/<u>E</u>35A in the mature sequence;
Δ9/13: NGF mutant protein with deletion of N-terminal 9-13 residues in the mature sequence;
DMEM: Dulbecco’s Modified Eagle’s Medium;
PMSF: phenylmethylsulfonyl fluoride;
SDS-PAGE: sodium dodecyl sulfate polyacrylamide gel electrophoresis;
DMSO: dimethyl sulfoxide;
AS: anti-sense oligos;
MM: mis-matched oligos;
E15.5 DRG: embryonic day 15.5 dorsal root ganglion;
ECD: extracellular domain;
MEM: Minimum Essential Medium;
TIRF: Total Internal Reflection Fluorescence;
EMCCD: electron multiplying charge-coupled device;
PC12: a cell line derived from a pheochromocytoma of the rat adrenal medulla;
BSA: Bovine Serum Albumin;
PBS: phosphate buffered saline;
IgG: Immunoglobulin G;
DIC: Days In Culture;

## Introduction

Nerve growth factor (NGF), discovered as a result of potent trophic actions on sensory and sympathetic neurons of the peripheral nervous system (PNS) in the 1950s(Levi-Montalcini and Hamburger, 1951), also regulates the trophic status of striatal and basal forebrain cholinergic neurons (BFCNs) of the central nervous system (Levi-Montalcini and Hamburger, 1951; Svendsen et al., 1994; Li and Jope, 1995; Kew et al., 1996; Conover and Yancopoulos, 1997; Lehmann et al., 1999). With the discovery of brain-derived neurotrophic factor (BDNF), Neurotrophin 3 (NT-3) and Neurotrophin 4 (NT-4), NGF is now known as a member of the neurotrophin family(Chao and Hempstead, 1995; Huang and Reichardt, 2001; Chao, 2003). NGF acts through two known receptors, the 140 kD tyrosine receptor kinase A (TrkA) and the 75 kD neurotrophin receptor (p75^NTR^) to transmit signals to the cytoplasm and nucleus of responsive neurons (Bothwell, 1995; Chao and Hempstead, 1995; Kaplan and Miller, 1997). NGF signaling through TrkA elicits many of the classical neurotrophic actions ascribed to NGF(Loeb and Greene, 1993). TrkB and TrkC mediate the signaling of NT-3 and NT-4, respectively(Huang and Reichardt, 2001; Chao, 2003). NGF as well as all members of the family also signal through p75^NTR^(Huang and Reichardt, 2001; Chao, 2003). p75^NTR^ contributes to sphingomyelin-ceramide metabolism (Dobrowsky et al., 1994; Dobrowsky et al., 1995) and modulates RhoA activity to regulate axonal growth(Yamashita et al., 1999; Gehler et al., 2004). In addition, p75^NTR^ has been shown to activate the NF-κB, Akt, and JNK pathways (Harrington et al., 2002; Roux and Barker, 2002) to either induce apoptosis or to promote cell survival and differentiation (Chao and Hempstead, 1995; Casaccia-Bonnefil et al., 1998; Salehi et al., 2000; Roux and Barker, 2002; Nykjaer et al., 2005).

Given its robust trophic effects, NGF has been investigated for therapeutic properties in neurodegenerative disorders(Apfel et al., 1994; Andreev et al., 1995; Apfel and Kessler, 1995; Blesch and Tuszynski, 1995; Anand et al., 1996; Apfel and Kessler, 1996; Apfel et al., 1998; Apfel, 1999b, a, 2000, 2002; Aloe et al., 2012). In one example, NGF’s robust trophic effects on BFCNs has suggested a role in treating AD in which this population degenerates (Olson, 1993; Hefti, 1994; Scott and Crutcher, 1994; Blesch and Tuszynski, 1995; Knusel and Gao, 1996; Koliatsos, 1996; Eriksdotter Jonhagen et al., 1998; Williams et al., 2006; Mufson et al., 2008; Schindowski et al., 2008; Schulte-Herbruggen et al., 2008; Cuello et al., 2010). Unfortunately, features of the biology of NGF have limited the extent to which it could be evaluated. Inability to cross the blood brain barrier prevented systemic administration. Delivery via the ventricular system even at low doses resulted in pain and studies in primates demonstrated Schwann cell hyperplasia that served to compromise CSF flow (Winkler et al., 1997). A recent Phase II trial in which NGF was delivered via virus to the basal forebrain demonstrated safety and was without pain(Tuszynski et al., 2015). Marked sprouting of BFCN fibers was evidence of a potent trophic effect but cognitive measures were unaffected(Rafii et al., 2014). Significant efforts were also invested to investigate NGF as a treatment for diabetic polyneuropathy(Apfel and Kessler, 1995; Anand et al., 1996; Apfel and Kessler, 1996; Tomlinson et al., 1996; Elias et al., 1998; Apfel, 1999b; Goss et al., 2002; Murakawa et al., 2002; Kanda, 2009). NGF treatment demonstrated some benefit in Phase II trial at 0.1 and 0.3μg/kg, but associated with dose-dependent hyperalgesia at the injection site(Apfel, 1999b, a). A large scale Phase III trial with a dose of 0.1μg/kg showed no beneficial effect (Apfel et al., 1998; Apfel, 2002).

NGF is not only a trophic factor but also functions as one of the key molecules for mediating inflammatory pain and neuropathic pain in the PNS (Lewin and Mendell, 1993; Lewin et al., 1993; Chuang et al., 2001; Watanabe et al., 2008). As such, clinical trials in which large doses of NGF were infused in patients with AD had to be terminated due to the extreme side effects of pain(Aloe et al., 2012). Other clinical trials using NGF in treating diabetic neuropathies and peripheral neuropathies in HIV were also discontinued after reports of serious side effects such as back pain, injection site hyperalgesia, myalgia, and weight loss (Hellweg and Hartung, 1990; Lein, 1995; Apfel et al., 1998; Unger et al., 1998; Rask, 1999; McArthur et al., 2000; Quasthoff and Hartung, 2001; Schifitto et al., 2001; Apfel, 2002; Pradat, 2003; Walwyn et al., 2006). Thus, the adverse effect of significant pain caused by NGF has severely limited its therapeutic use in treating neurodegenerative disorders. To overcome these pain-causing side effects of NGF, it is of paramount importance to elucidate the role of NGF and its receptor signaling by TrkA and p75^NTR^ in nociception.

A large body of genetic and clinical evidence has pointed to both TrkA and p75^NTR^ as contributing factors to sensitization of inflammatory pain mediated by NGF. For example, recessive mutations in TrkA cause hereditary sensory and autonomic neuropathy type IV (HSAN IV) (OMIM # 256800), also known as congenital insensitivity to pain with anhidrosis (Indo, 2001, 2002). Strong evidence supports a role of TrkA in mediating the sensitization effect of NGF: attenuation of TrkA expression(Malik-Hall et al., 2005; Alvarez and Levine, 2014) and pharmacological inhibition of TrkA-mediated signaling pathways extracellular signal-related kinase (ERK), phosphatidylinositol 3-kinase (PI3K), and phospholipase Cγ (PLCγ) all reduced NGF-induced hyperalgesia (Fang et al., 2005; Malik-Hall et al., 2005; Summer et al., 2006; Mantyh et al., 2011; Alvarez and Levine, 2014; Ashraf et al., 2016). Furthermore, NGF still evoked hyperalgesia in mice lacking p75^NTR^, pointing to the involvement of TrkA(Bergmann et al., 1998). Evidence that p75^NTR^ has a role in pain signaling pathways is largely indirect. In one example, injecting a neutralizing antibody to p75^NTR^ prevented NGF-induced pain behavior and NGF-mediated increases in action potentials in sensory neurons (Zhang and Nicol, 2004; Watanabe et al., 2008; Iwakura et al., 2010). Therefore, both TrkA and p75^NTR^ signals contribute to pain induced by NGF. However, how these two receptor(s) interact to mediate pain is poorly defined.

Recently, patients in consanguineous Swedish families suffering from length-dependent loss of pain that often leads to bone fractures and joint destruction were shown to harbor a homozygous missense mutation in NGF(Einarsdottir et al., 2004; Carvalho et al., 2011). The disorder was labeled hereditary sensory autonomic neuropathy type V (HSAN V) (Online Mendelian Inheritance in Man (OMIM) # 608654). Genetic analysis of these HSAN V patients revealed a point mutation (661C>T) causing a substitution of tryptophan (W) for arginine (R) at position 211 in the pro-form of the NGF polypeptide (pro-NGF^R221W^); this residue corresponds to the position 100 in the mature protein (NGF^R100W^) (Einarsdottir et al., 2004). Unlike HSAN type IV, which results from mutations in TrkA, HSAN V patients appear to have normal cognitive function, suggesting that the mutant NGF may retain its trophic functions in the CNS (Einarsdottir et al., 2004). We and others reasoned that NGF^R100W^ provides a tool to decipher possible differences in the trophic and nociceptive actions of NGF (Covaceuszach et al., 2010; Capsoni et al., 2011; Capsoni, 2014).

Initial characterization of NGF^R100W^ revealed that the R100 mutation may disrupt the processing of pro-NGF to mature NGF in cultured cells, resulting in relatively higher percentage of NGF secreted as the pro-form (Larsson et al., 2009). Given the difficulties in expressing NGF^R100W^, Catanneo and colleagues examined a different series of residues at position 100, including NGF^R100E^ and NGF double mutant (NGF^P61S/R100E^) using recombinant techniques(Capsoni et al., 2011). They discovered that these NGF^R100^ mutants bound normally to TrkA, but failed to bind to p75^NTR^ (Covaceuszach et al., 2010). This finding suggested that failure to activate p75^NTR^ signaling was sufficient to attenuate pain induced by NGF and suggesting that TrkA signaling had little or no effect in pain(Capsoni et al., 2011). The authors speculated that NGF^R100^ mutant would allow for development of p75^NTR^ antagonists, such as NGF^R100W^ as a ‘painless NGF’ therapeutic agent (Malerba et al., 2015).

In the present study, we examined the binding and signaling properties of the mature form of the naturally occurring mutant NGF (NGF^R100W^) in HSAN V and compared the effects to those of wtNGF and NGF mutants that selectively engage and signal through either TrkA or p75^NTR^. We discovered that NGF^R100W^ retains binding and signaling through TrkA to induce trophic effects but not binding or activation of p75^NTR^. Our findings are evidence that NGF^WT^ acts through both p75^NTR^ and TrkA to cause pain. Our findings are consistent with a necessary role for TrkA in both acute sensitization and delayed priming of nociceptive responses. In contrast, signaling through p75 appears to be unnecessary for acute sensitization but does contribute to priming. The integration of both TrkA and p75 signaling thus appears to regulate neuroplastic effects of NGF in peripheral nociception.

## Materials and Methods

### Ethic Statement

All experiments involving the use of animals were approved by the Institutional Animal Care and Use Committee of University of California San Diego and University of California San Francisco. Surgical and animal procedures were carried out strictly following the NIH Guide for the Care and Use of Laboratory Animals.

### Chemicals, oligos and reagents

Streptavidin-QD605, −655 (Q10103MP, Q10123MP) conjugates were from Invitrogen, all other chemicals were from Sigma unless noted otherwise. Recombinant extracellular domains of p75^NTR^ were a generous gift from Dr. Sung Ok Yoon of Ohio State University. Prostaglandin E2 (PGE_2_) was purchased from Sigma (Cat# 82475). NGF (wtNGF, NGF^R100W^, KKE, Δ9/13) proteins were produced in our own laboratory(Sung et al., 2011).

### Cloning

Mouse pro-NGF was amplified by PCR from a NGF-GFP plasmid (a generous gift from Professor Lessmann, Mainz, Germany). The forward primer sequence was: 5’-acgaattccaccatgtccatgttgttctacactctgatcactgcg-3’ and the reverse primer sequence was: 5’-gatggatccttcgtgccattcgattttctgagcctcgaagatgtcgttcagaccgccaccgacctccacggcggtggc-3’. The reverse primer contains a sequence coding for the 17-amino acid AviTag: GGGLNDIFEAQKIEWHE. The sequence was based on the #85 AviTag peptide sequence described previously (Schatz, 1993). One glutamic acid residue was added to the C-terminal AviTag based on a finding by Avidity (http://www.avidity.com) that it greatly enhanced the biotinylation rate of the AviTag (Beckett et al., 1999). Platinum pfx DNA polymerase (Invitrogen, Cat#11708021) was used following the manufacture’s instruction. The 50 μl of reaction was denatured at 94°C for 4 min, followed by 25 cycles of amplification (30 s at 94°C; 30 s at 50°C; 90 s at 68°C). An additional extension was performed at 68°C for 4 min. The PCR product was purified and digested with EcoRI (Fermentas, Cat# FD0274) and BamHI (Fermentas, Cat# FD0054). It was ligated in-frame into the pcDNA3.1-myc-His vector that was predigested with EcoRI/BamHI. The resulting construct was designated as pcDNA3.1-NGFavi. BirA was amplified by PCR from pET21a-BirA (purchased from http://www.Addgene.com; Plasmid# 20857) (Howarth et al., 2005) using a forward primer (forward primer: 5’-gtgaac atg gctagcatgact-3’) and a reverse primer (5’-ggtgctcgagtcatgcggccgcaagct-3’ (containing an XhoI site). The PCR was carried out using Pfx as described above. The PCR product was digested with XhoI (Fermentas, Cat# FD0694) and was subcloned into pcDNA3.1 myc.his (+) vector (Invitrogen) that was pre-cut with EcoRV (Fermentas, Cat# FD0303) and XhoI. The resulting plasmid was designated as pcDNA3.1-BirA. All primers were purchased from Elim Biopharmaceuticals, Inc (Hayward, CA). All constructs were verified by sequencing (Elim Biopharmaceuticals, Inc.). NGF^R100W^ was cloned using the same method, but with a point mutation at 661 (C>T). The KKE and Δ9/13 mutant constructs were obtained from Dr. K. Neet of Rosalind Franklin University (Hughes et al., 2001; Mahapatra et al., 2009), were subcloned to the pcDNA3.1-myc-His vector with an Avitag (Sung et al., 2011).

### Protein purification

HEK293FT cells were grown in 15 cm plates to 70% confluency. Cells were changed to 25 ml of DMEM-high glucose, serum-free media that was supplemented with 50 μM d-biotin (Sigma, Cat# B4639). 15-21 μg pcDNA3.1-NGFavi, NGF^R100W^avi, KKEavi, and Δ9/13avi plasmids DNA plus 15-21 μg pcDNA3.1-BirA plasmids DNA were mixed with 1 ml of DMEM-high glucose media and 60 μl of Turbofect (Fermentas, Cat# R0531). The mixture was incubated at room temperature for 15 min and was then added into the media by drop-wise method. Transfected HEK293FT cells were incubated at 37°C, 5% CO_2_. 72 hrs post transfection, media were collected for protein purification.

Media were harvested and adjusted to 30 mM phosphate buffer, pH 8.0, 500 mM NaCl, 20 mM imidazole, and a cocktail of protease inhibitors (1 mM PMSF from Sigma Cat# P7626, and 1 μl/ml aprotinin from Sigma, Cat#A6279). After incubation on ice for 15 min, media were cleared by centrifugation at 18,000 rpm for 30 min using a Beckman JA-20 rotor. Ni-NTA resins (Qiagen, Cat# 30250) were rinsed with the washing buffer (30 mM phosphate buffer, pH8.0, 500 mM NaCl, 20 mM imidazole, and a cocktail of protease inhibitors from Sigma, Cat# S8820). Ni-NTA resins were added to the media at a concentration of 0.3 ml Ni-NTA/100 ml media and incubated overnight with rotation at 4°C. The media/Ni-NTA slurry was loaded onto a column and the captured Ni-NTA resins were washed with 10 ml wash buffer and eluted with 5 ml elution buffer (30 mM phosphate buffer, pH8.0, 500 mM NaCl, 300 mM imidazole, protease inhibitors). Every 500 μl volume of elution was collected. The purity and concentration of NGF was assessed by SDS-PAGE using a silver staining kit (Fast silver, G-Biosciences, Cat# 786-30, Maryland Heights, MO). Known quantities of NGF purified from mouse sub maxillary glands were used as standards. The first two eluted fraction normally contained most of purified proteins.

### Cell culture and transfection

PC12 cells or a subclone of PC12 cells, PC12M, PC12^nnr5^ cells were cultured as described previously (Wu et al., 2001; Wu et al., 2007). NIH3T3-TrkA, NIH3T3-p75^NTR^ cells were as described (Huang et al., 1999). HEK293FT cells (Invitrogen, Cat# R70007) cells were cultured in DMEM-high glucose media (4.5 g/l glucose, Mediatech, Cat#10-013-CV), 10% FBS, 1% penicillin/streptomycin.

### Administration of wtNGF, NGF^R100W^, prostaglandin E2 (PGE_2_), oligos and inhibitors to adult rats

Experiments were performed on adult male Sprague-Dawley rats (220-240g, Charles River, Hollister, CA, USA). All experiments were carried out following protocols that have been approved by the University of California San Francisco Committee on Animal Research and conformed to National Institutes of Health Guidelines for the Care and Use of Laboratory Animals. Either 200 ng of wtNGF or NGF^R100W^ was injected intradermally on the dorsum, of one hind paw of adult rats. Nociceptive threshold in the injected paws were then tested over time.

For studies using K252a (Sigma#: K1639) and GW4869 (Sigma#: D1692), both inhibitors were dissolved in DMSO (2 μg/μl) and then diluted to a concentration of 0.2 μg/μl in saline at the time of the experiments. Five mins prior to injection of NGF, 5 μl (1 μg) of inhibitors were administered intradermally on the dorsum of the hind paw, at the same site at which NGF was injected. The experimental design is illustrated in Figure 7A.

### Randall-Selitto mechanical and Hargreaves thermal tests

Mechanical nociceptive threshold measured using the Randall-Selitto paw pressure test (Randall and Selitto, 1957) with an Ugo Basile Algesymeter (Stoelting, Chicago, IL, USA) (Ferrari et al., 2010; Ferrari et al., 2013; Ferrari et al., 2015a; Ferrari et al., 2015b). This device exerts a linear increase in force to the dorsum of the hind paw of the rats. Prior to the test, rats were kept in individual restrainers for 20 min to acclimatize them to the experimental environment. The restrainers had openings that allowed rats to extend hind paws during the test. Mechanical thresholds were calculated as the average of three readings.

Thermal threshold was measured using the Hargreaves test, which applies heat stimuli by an adjustable high-intensity movable halogen projector lamp (Malmberg and Yaksh, 1993). Baseline responses were first measured and were averaged from at least two readings. 600 ng of wtNGF or NGF^R100W^ was then injected in the plantar surface of one hind paw, under isofluorane anesthesia. After recovery from the anesthesia, in less than a min, they were placed on a glass plate, and acclimatized for 20 min. Thermal nociceptive thresholds were evaluated within 1 hr after the injections.

### Binding and internalization assays of NGF-QD

At 50% confluence of NIH3T3-TrkA and NIH3T3-p75^NTR^, cells were starved in serum free DMEM-high glucose media for 4 h at 37°C. NGF or NGF mutants were conjugated with QD 655-streptavidin (Invitrogen, Cat# Q10121MP) on ice 4°C for 30 min. 0.2 nM of conjugate was applied to cells. Cells were incubated at 20°C for 20-30 min, washed with serum free DMEM-high glucose media, and then surface binding quantified. For the internalization assay, cells were incubated with 1nM conjugates for 30 min or 2 h, for 3T3-TrkA and 3T3-p75^NTR^ respectively at 37°C. Cells were washed, then subjected to imaging.

### Pull down assay

The recombinant protein, Fc-75^NTR^ ECD (extracellular domain) was a kind gift from Dr. Sung Ok Yoon, the Ohio State University. 5 μl of supernant from insect lysate expressing Fc-p75^NTR^ was incubated with a range of either wtNGF or NGF^R100W^ (0~10ng) for overnight at 4°C. Avidin-agarose beads (20 μl) were added/incubated for 2 hrs at 4°C. Beads were washed, boiled with protein sample buffer and subjected to SDS-PAGE. Western blotting was performed using the function blocking antibodies against the extracellular domain of p75^NTR^ (REX) (Mischel et al., 2001).

### DRG culture, live cell imaging and data analysis

Embryonic dorsal root ganglions (DRGs E15-16) were isolated from Sprague-Dawley as described in (Cui et al., 2007; Wu et al., 2007; Sung et al., 2011) with some minor modifications. Cells were maintained with alternation between growth media (MEM media containing 10% heat inactivated FBS and 100 ng/ml of NGF) and selection media (MEM media containing 0.5-1 gM of cytosine β-d-arabinofuranoside (Sigma: C1768), and 100 ng/ml NGF) every two days.

For survival analysis, only the cells with round and transparent cell bodies were counted as DRG neurons to exclude possible fibroblasts populations. In phase-contrast image, the cell bodies of DRGs look whitish and transparent whereas fibroblasts have black and flattened shaped cell bodies.

For live cell imaging, dissociated DRGs were cultured in microfluidic chambers for 7–10 days (Cui et al., 2007). The microfluidic chambers, manufactured in house, were plated onto 24 mm × 48 mm glass coverslips that were pre-coated with poly-l-lysine (Sigma, P8920) as described in (Taylor et al., 2006). Dissociated DRG neurons were plated into the cell body chamber. The growth/selection scheme outlined above was repeated. Axons from the DRG neurons started to cross the microgrooves after 3 days and reached the axonal chamber in another 7–8 days. Prior to live imaging of axonal transport of NGF, all compartments (cell body and axonal chambers) of the DRG neurons were thoroughly rinsed and were depleted of NGF in NGF-free, serum-free MEM media for 3 h. NGF-QD605 was prepared following the protocol described above. NGF-QD605 was added to a final concentration of 0.2 nM to the axonal chambers for 2 h at 37°C. Live cell imaging of NGF-QD605 transport with the axons was carried out using a modified inverted microscope (Nikon TE300) for pseudo-TIRF illumination (Zhang et al., 2010). The microscope stage was equipped to maintain a constant temperature (37°C). CO_2_ level (5%) was maintained by using CO_2_ independent medium (Invitrogen, #18045-088) during live imaging. The laser beam of 532 nm was used and penetrated ~ 1 μm into aqueous solution at an incident angle. Fluorescence emission was filtered with QD605/20 emission filter (Chroma Technology, Rockingham, VT). Time-lapse images were acquired at the speed of 10 frames/s and were captured using an EMCCD camera (Cascade 512B, Photometric, Tucson, AZ). All data was processed and analyzed using a MATLAB software pipeline.

**Single Cell Patch Clamp**: All recordings were obtained from small to medium diameter sized cells from cultured DRGs at room temperature. One electrode whole-cell voltage-clamp recording was performed using Axopatch-1D amplifiers (Axon Instruments). 2-5 MW sized patch electrode and puffing pipette were used. All cells were clamped at −60 mV holding potential to measure low pH-evoked current.

Standard external solution containing 145mM NaCl, 5mM KCl, 2mM CaCl_2_, 1mM MgCl_2_, 10mM HEPES, and 10 mM glucose (adjusted to pH 7.4 with NaOH) was used to incubate DRG neurons. To block possible synaptic transmission via activation of ionotropic receptors, 1mM TTX, 10mM CNQX, 50mM AP5, and 50mM picrotoxin were added to the bath. Internal recording solution contained 130mM K-gluconate, 10mM HEPES, 0.6mM EGTA, 5mM KCl, and 2.5mM Mg-ATP (adjusted to pH 7.3 with KOH). Low pH puffing solution was made from external solution and was adjusted to pH 5.5 by adding 1N HCl.

Cells were starved for 2 hrs with MEM prior to recording. To evoke low pH response, pH 5.5 external solution was applied briefly through an additional glass pipette placed 50 mm away from the recorded cell and by a Picospritzer (5-10 psi, for 50-100 ms, Picospritzer II, General Valve). The responses seen at this configuration were not evoked by mechanical stimulation due to air puffing as when neutral pH solution was used, so no response was observed. wtNGF or NGF^R100W^, at a concentration of 50 ng/ml was applied directly to the bath and incubated for 10 min followed by low pH puffing. Data analysis was done in pClamp (Molecular devices) by calculating the charge transfer for 100 ms and normalized by computing the ratio of response after NGF/response before NGF.

### Antibodies and SDS-PAGE/blotting

Standard protocols were followed for SDS-PAGE and blotting. Rabbit anti-pro-NGF IgGs were a generous gift of Dr. B. L. Hempstead of Cornell University. Rabbit anti-NGF IgGs were from Santa Cruz Biotechnology Inc (Santa Cruz, CA, Cat# sc-549). Rabbit IgGs against Trk-pTyr490, pErk1/2, total Erk1/2 and p-Cofilin were from Cell Signaling Technology (Danvers, MA, Cat #9141, 9101, 9102, 3311 respectively). Mouse IgGs against pJNK, mouse IgGs against total PLC-γ were from Santa Cruz (sc-6254, sc-7290 respectively), and rabbit IgGs against pPLC-γ were from GeneTex (GTX61714). Rabbit functional blocking antibodies against the extracellular domain of p75^NTR^ (REX) (Mischel et al., 2001) was a generous gift of Dr. L. Reichardt of UCSF. Rabbit monoclonal antibodies against pAkt were from Epitomics Inc (Cat#2214-1). Rabbit anti-AviTag IgGs were purchased from Genescript (Piscataway, NJ, Cat# A00674).

### Immunostaining

Immunostaining was carried out according published protocols(Weissmiller et al., 2015). Briefly, PC12^nnr5^ cells were cultured on coverslips that were pre-coated with Matrix-gel (BD). Following serum starvation for 2hr, cells were treated with either 50 ng/ml wtNGF or 50 ng/ml NGF^R100W^. Cells were then fixed for 10 mins with 4% paraformaldehyde at 37°C and permeabilized for 15°C at room temperature with 0.1% triton X-100. 3% bovine serum albumin (BSA) and 5% goat serum in PBS was used for blocking. Then the cells were incubated for overnight at 4°C with blocking solution containing 1/100 diluted primary antibody, active RhoA from NewEast Biosciences (Cat# 26904). After washing three times primary antibody with PBS, and rocking 5 min, the cells were incubated with a 1/800 dilution of Alexa Fluor 488-goat anti-mouse IgG (from Molecular Probes Cat# A1100) for an hour at room temperature with rocking covered with foil. After three washes with PBS, nuclei were labeled with 1 μg/ml Hoechst 33342 (bisBenzimide H 33342 trihydrochloride, Sigma Cat#B2261) for 5 min at room temperature. Cells were rinsed, air-dried, mounted and examined with a Leica microscopy using a 100x oil objective lens.

### Statistical analysis

All experiments were repeated at least three times, independently. Statistical analyses of results and calculation of *p* values were performed using Prism5 (GraphPad Software, La Jolla, CA). For un-pairwise comparisons, the Student’s t-test was used. For multiple comparisons, the Tukey one way ANOVA test was used.

## Results

### 1. NGF^R100W^ does not elicit acute thermal or mechanical hyperalgesia *in vivo*

Previous studies suggested that the NGF mutation associated with HSAN V disrupted either the processing of the pro-form (i.e. NGF^R221W^) or secretion of the mature form (NGF^R100W^), resulting in preferential secretion of the NGF^R221W^ (Larsson et al., 2009; Covaceuszach et al., 2010; Carvalho et al., 2011; Capsoni, 2014). However, these findings and interpretations are perplexing since HSAN V patients: 1) report no autonomic symptoms (Capsoni, 2014), which would argues against increased levels of pro-NGF, a p75 ligand that induces death of sympathetic neurons(Roux and Barker, 2002; Nykjaer et al., 2005; Khodorova et al., 2013); and 2) have intact mental abilities, arguing against clinically meaningful CNS neuronal loss and dysfunction. To further explore the binding, signaling and actions of NGF^R100W^, we produced and characterized the mature form of the protein in HEK293 cells.

While superficial sensation in patients homozygous for NGF^R100W^ is normal, it is unknown if this mutant NGF can induce nociceptor sensitization or transition to chronic pain. Therefore, we measured the behavioral response of adult rats to mechanical stimuli or noxious thermal stimulus after a single injection of either wtNGF or NGF^R100W^. Proteins were administered intradermally to the hind paws of adult male Sprague-Dawley rats following published protocols (Taiwo et al., 1991).

We used the Randall-Sellito method to measure mechanical threshold (Randall and Selitto, 1957). For all experiments, baseline values were measured before the injection of NGF. Previous Reports showed that 10ng-1μg of injected wtNGF induced significant mechanical hyperalgesia (Andreev et al., 1995; Malik-Hall et al., 2005). When 200 ng of wtNGF (**Fig 1A, B, C** blue) was injected to the dorsum of the rat’s hind paw, the mechanical nociceptive threshold was reduced ~25 % compared to baseline (p<0.0001, unpaired t-test, 0m vs 60m/1^st^ day) (**Fig 1B, C** blue). Mechanical hyperalgesia was evident as soon as 15 min and reached the maximum by 1 h, an effect that lasted for ≥24 h. Hyperalgesia was diminished to ~15% by 5 days (**Fig 1C**). In marked contrast to wtNGF, injection of 200 ng of NGF^R100W^ had no effect on mechanical nociceptive threshold (**Fig 1B, C** red).

**Figure 1.**
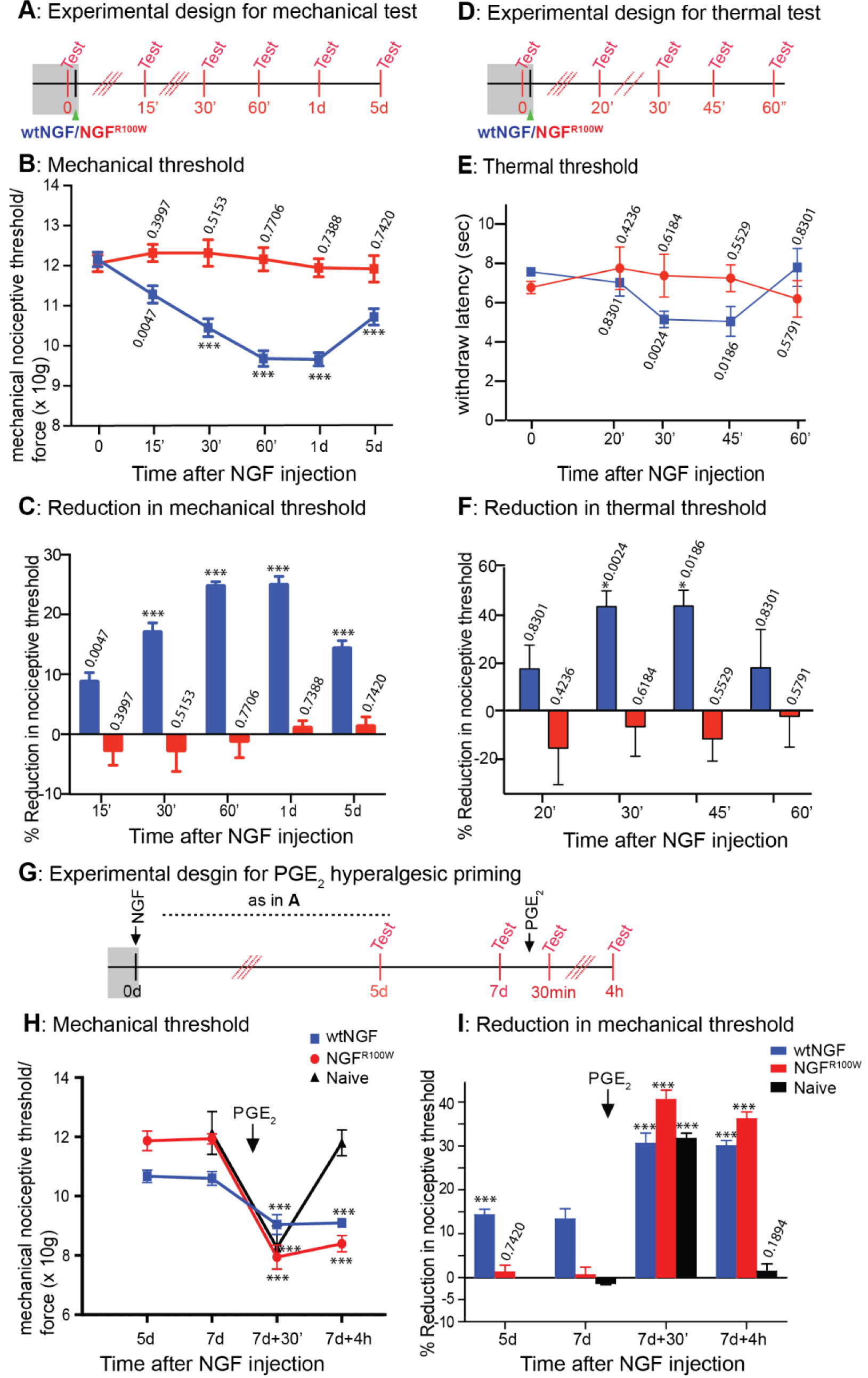
NGF^R100W^ does not induce acute sensitization in adult rats. **Mechanical hyperalgesia (A, B, C)**. The Randall-Selitto method was used to measure mechanical hyperalgesia in adult rats. Following intradermal injection of 200 ng of wtNGF (n=6) or NGF^R100W^ (n=6) into the rat’s hind paws mechanical threshold was measured at indicated times. A significant decrease in mechanical nociceptive threshold in rats injected with wtNGF was seen within 15 min, maximum by 1 hr and of duration at least 5 days. In contrast, NGF^R100W^ did not produce a significant decrease in the threshold during the 5 days period. **Thermal hyperalgesia (D, E, F)**. The Hargreaves test was used to measure thermal hyperalgesia. Baseline response was first measured for each test animals before NGF injection. Hindpaws were injected with 600 ng of either wtNGF or NGF^R100W^. Thermal threshold was measured at indicated times. Animals that were injected with wtNGF showed a significant decrease at 30 min. The decrease was dissipated after an hour. NGF^R100W^ failed to reduce thermal nociceptive threshold during the entire 1-hour test period. Six rats for each group (n=6) were used in the test. Unpaired t-test were performed against the baselines within each NGF injection group to produce p-values. Comparions were done between the threshold before injection with either wtNGF or NGF^R100W^ to produce p-values that were noted in the figure. *** indicates when p<0.001 **PGE_2_-hyperalgesic priming effect (G, H, I)**. On the 7^th^ day after NGF administration, PGE_2_ (prostaglandin E_2_) was injected intradermaly (100 ng/5μl) to test hyperalgesic priming. Naïve rats (non-treated with NGF) are shown as a control. Naïve animals showed acute hyperalgesia within 30 min, but not at 4hrs, after PGE_2_ injection. Rats pretreated with intradermal injection of wtNGF showed decrease in mechanical nociceptive threshold after PGE_2_ injection at 30 min, like the controls, but which lasted longer than 4 hr in contrast to naïve animal. Rats pretreated with NGF^R100W^ showed significant decrease in mechanical nociception after PGE_2_ injection, which lasted at least 4 hours comparably to wtNGF. Data are presented as mean ± SEM. Unpaired t-tests were performed versus values at ‘before PGE_2_ injection’. *** indicates when p<0.001 Six paws (n=6) were used for wtNGF or NGF^R100W^.

We then measured thermal hyperalgesia using the Hargreaves method (Hargreaves et al., 1988). Intraplantar injection of 600 ng of wtNGF to the hind paw of rats induced thermal hyperalgesia, as demonstrated by as much as a 34.8 % decrease in nociceptive threshold (p=0.0186, 0m vs 45m, unpaired t-test) (**Fig 1D, E, F** blue). Acute thermal hypergelsia was first observed at 20-30 min after injection with the maximal effect at 45 min followed by a return to base line (**Fig 1E, F** blue). In contrast, injection of 600 ng of NGF^R100W^ did not induce acute thermal hyperalgesia (**Fig 1E, F** red). We conclude that, unlike wtNGF, NGF^R100W^ does not induce acute thermal or mechanical hyperalgesia.

### 2. NGF^R100W^ induced hyperalgesic priming

To test if NGF^R100W^ could contribute to chronic pain, we took advantage of the hyperalegesic priming paradigm with the injection of prostaglandin E2 (PGE_2_) (Reichling and Levine, 2009). Because wtNGF was shown to induce hyperalgesic priming in rats (Ferrari et al., 2010), we used this model to examine responses to wtNGF and NGF^R100W^. 7 days post injection of 200ng of either wtNGF or NGF^R100W^, PGE_2_ (100 ng/5μl) was injected into the same injection site into adult rats (**Fig 1G**). If a priming effect for chronic pain is absent, injection of PGE_2_ only induces acute hyperalgesia that disappears by 4hrs (Ferrari et al., 2010; Ferrari et al., 2013; Ferrari et al., 2015b). However, in the presence of priming, the same dose of PGE_2_ induces a much prolonged hyperalgesia to mechanical stimuli (Aley and Levine, 1999; Reichling and Levine, 2009).

As previously reported (Ferrari et al., 2010), intradermal injection of wtNGF (200ng) induced acute mechanical hyperalgesia that lasted through day 7 (**Fig1H, I**). As before, NGF^R100W^ failed to induce acute mechanical hyperalgesia (**Fig 1H, I**). On the 7^th^ day after measuring the threshold baseline, PGE_2_ was injected into the same site as for NGF and the mechanical threshold was measured at 30 min and 4 hr thereafter. Consistent with previous findings (Aley and Levine, 1999; Ferrari et al., 2010), in animals treated with wtNGF, PGE_2_ induced prolonged hyperalgesia unattenuated at least for 4 hrs (**Fig 1H, I** blue), while in naïve subjects, PGE_2_ induced only acute hyperalgesia at 30 min and the hyperalgesic effect was largely dissipated at 4 hrs (**Fig 1 H, I** black, naïve). Remarkably, NGF^R100W^ was also as potent as wtNGF in inducing priming with prolonged PGE_2_ hyperalgesia, lasting at least 4 hrs (**Fig 1H, I** red, NGF^R100W^). We conclude that NGF^R100W^ retains binding and signaling necessary to induce hyperalgesic priming (**Fig 1 H, I**), despite of its inability to induce acute hyperalgesia (**Fig 1B,C,E,F**). These findings suggest that studies of NGF^R100W^ may provide insights into mechanisms underlying the nociceptive and trophic functions of NGF in sensory neurons.

### 3. NGF^R100W^ does not potentiate low H^+^-evoked response in sensory neurons *in vitro*

Among the known mechanisms of pain transduction, NGF produces acute hypersensitivity by potentiating nociceptive ion channels such as the capsaicin receptor (also known as TRPV1) and acid-sensing ion channels (ASICs) (Qiu et al.; McCleskey and Gold, 1999; Szallasi and Blumberg, 1999; Chuang et al., 2001; Julius and Basbaum, 2001; Mamet et al., 2003; Yen et al., 2009). ASICs and TRPV1 are proton-gated non-selective cation channels that mediate acid evoked pain in peripheral sensory neurons (Caterina et al., 2000; Davis et al., 2000; Chen et al., 2002; Hellwig et al., 2004). We next performed patch clamp to test if NGF^R100W^ no longer elicited nociceptive response at the cellular level in cultured rat E15.5 dorsal root ganglion (DRG) sensory neurons. At DIC5 (<u>D</u>ays-<u>I</u>n-<u>Culture</u>), neurons were starved for NGF for 2 hrs. We used a moderate acidic solution (pH 5.5) to activate ASICs and TRPV1 by puffing the patch-clamped cell body (**Fig 2A**). Proton-evoked currents were measured before and after a 10 min application of either wtNGF or NGF^R100W^ (**Fig 2B**). The responses were normalized by calculating the ratio of the “after NGF” value:the “before NGF” value. Consistent with previous studies (Koplas et al., 1997) (Shu and Mendell, 1999a) (Shu and Mendell, 1999b), an 10 min of wtNGF treatment produced an acute sensitization, as displayed by a 1.5 fold increase in proton evoked current (**Fig 2C**). However, NGF^R100W^ did not induce hyper-sensitization (Fig 2C) (wtNGF vs NGF^R100W^; *p*<0.01; paired *t*-test).

**Figure 2.**
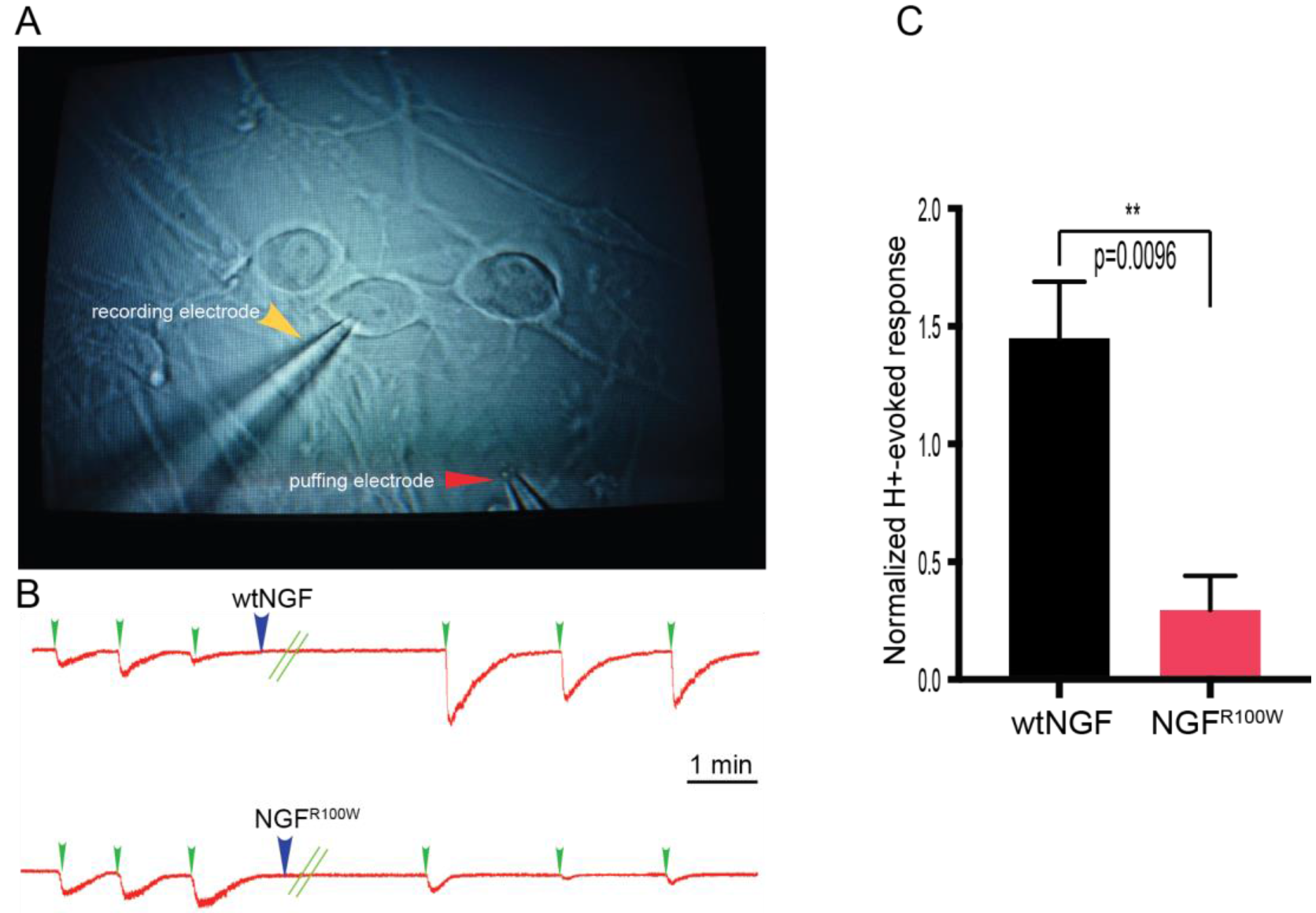
Low H^+^-evoked response by single cell patch clamping. Rat E15.5 DRG neurons were cultured as described in the Materials and Methods. At DIC5, DRG neurons were deprived of NGF for 2 hrs. In **A**, a phase-contrast image of DRG neurons (under 60x magnification) shows the experimental setup. The patch pipette approached from the left, while another glass pipette that applied brief puffs of pH 5.5 solution to the nearby cell body was placed at a distance. This pipette did not induce mechanical responses. **B**: Whole cell patch-clamp was performed at a holding potential of −60 mV in DRG. The proton-evoked response was measured following a brief application of moderate acidic solution of pH 5.5 (blue arrow) onto cell body to induce inward current (designated as “before NGF response”). 50 ng/ml of wtNGF or NGF^R100W^ (green arrow) was then applied to the bath solution for 10 min, three additional puffs were applied to record ‘after NGF’ response. The data was normalized by calculating ‘after NGF’ response/ ‘before NGF’ response and presented in **C**. Bar graphs represent mean ± SEM. wtNGF sensitized the inward current by 1.45 fold, but not NGF^R100W^ (p=0.0096; wtNGF versus NGF^R100W^; unpaired t-test).

### 4. NGF^R100W^ induces differentiation and supports survival of rat E15.5 DRG neurons

HSAN V patients show reduced responses to painful stimuli but retain normal cognitive function (Einarsdottir et al., 2004). We thus speculated that NGF^R100W^ would sustain trophic signaling resulting in the differentiation and survival of DRG sensory neurons (Winter et al., 1988; Wu et al., 2007). To compare the bioactivity of NGF^R100W^ with wtNGF, we performed a dose-response survival assay of DRG neurons for both wtNGF and NGF^R100W^ (0, 10, 50, 100ng/ml). We used E15.5 DRG and measured the number of healthy DRGs at Day 8 in culture after treating either with wtNGF or NGF^R100W^ (**Fig 3A**). At DIC8, Phase contrast images of DRG cultures were taken and survival were quantitated (**Fig 3B**). Unpaired t-test was performed to compare the number of healthy DRGs treated with wtNGF or NGF^R100W^. At any concentration of treatment, NGF^R100W^ exerted trophic effect to support the survival of DRGs comparably to wtNGF. As a control, cultures with no addition of NGF showed significant death (**Fig 3A**). We thus conclude that NGF^R100W^ supports the survival of rat E15.5 DRG neurons and that it does so as effectively as wtNGF.

**Figure 3.**
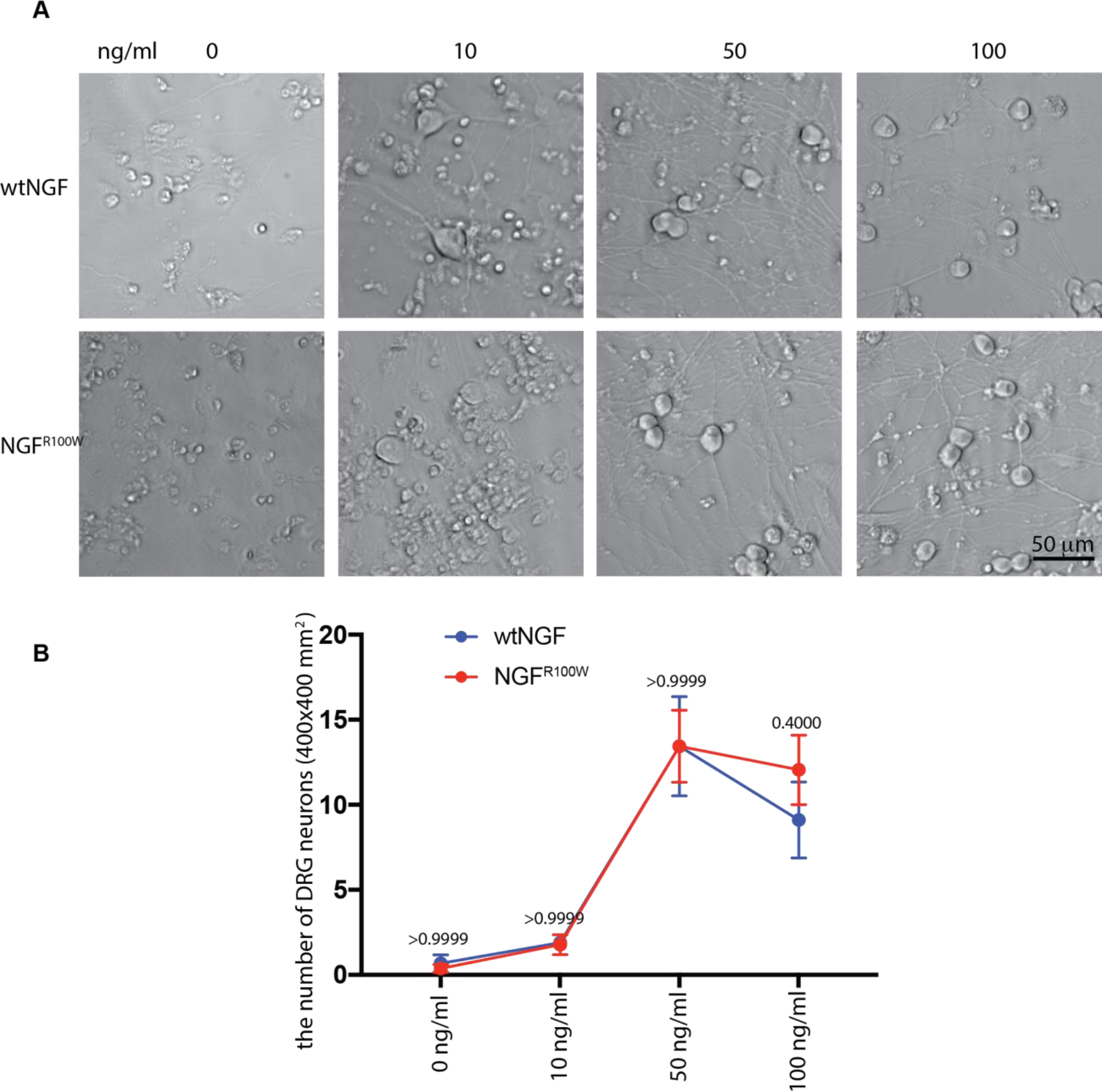
Dorsal root ganglion survival assays. Rat E15.5 DRG neurons were cultured as described in the Materials and Methods. Parallel cultures were supplied with either wtNGF oror NGF^R100W^ with range from 0 ng/ml to 100 ng/ml. Phase-contrast images of DRG neurons at DIV 8 were captured and representative images are shown (**A**). Negative control (no NGF, 0 ng/ml) failed to maintain the survival of DRGs. NGF^R100W^ maintained the survival as potently as wtNGF. (**B**) The survival rate of DRG neurons i.e. cell counts, by wtNGF or NGF^R100W^ was not significantly different. (**E**) Unpaired t-test was performed on wtNGF versus NGF^R100W^ (p>0.9999 at 0, 10, 50 ng/ml, p=0.4000 at 100 ng/ml).

### 5. NGF^R100W^ binds to and is internalized through TrkA, but p75^NTR^

NGF binds and signals through TrkA and/or p75^NTR^ receptors to effect neuronal function (Frade and Barde, 1998; Yoon et al., 1998; Sofroniew et al., 2001; Chao, 2003). We then explored if NGF^R100W^ differed from wtNGF in binding and internalization through TrkA and p75^NTR^. We used a NIH3T3 cell line that stably expresses either TrkA or p75^NTR^ (Hempstead et al., 1991; Kaplan et al., 1991; Zhou et al., 1994; Huang et al., 1999) to perform in-cell binding assays. NGF^R100W^ and wtNGF were each labeled with Quantum dots 655 (QD655) before incubating with either NIH3T3-TrkA-, or NIH3T3-p75^NTR^ cells. Saturable binding was demonstrated by employing a range of NGF concentrations (**Fig 4A, B**). Binding data were fit to a classical hyperbolic binding curve (one-site binding) using non-linear regression analysis using Prism 6. The results showed that NGF^R100W^ binding to TrkA (Kd = 2.79 nM) was essentially indistinguishable from that for wtNGF (Kd = 2.27 nM) (**Fig 4A**). In contrast, NGF^R100W^ showed minimal binding to p75^NTR^ even at concentrations at which wtNGF binding was essentially saturated (**Fig 4B**).

**Figure 4.**
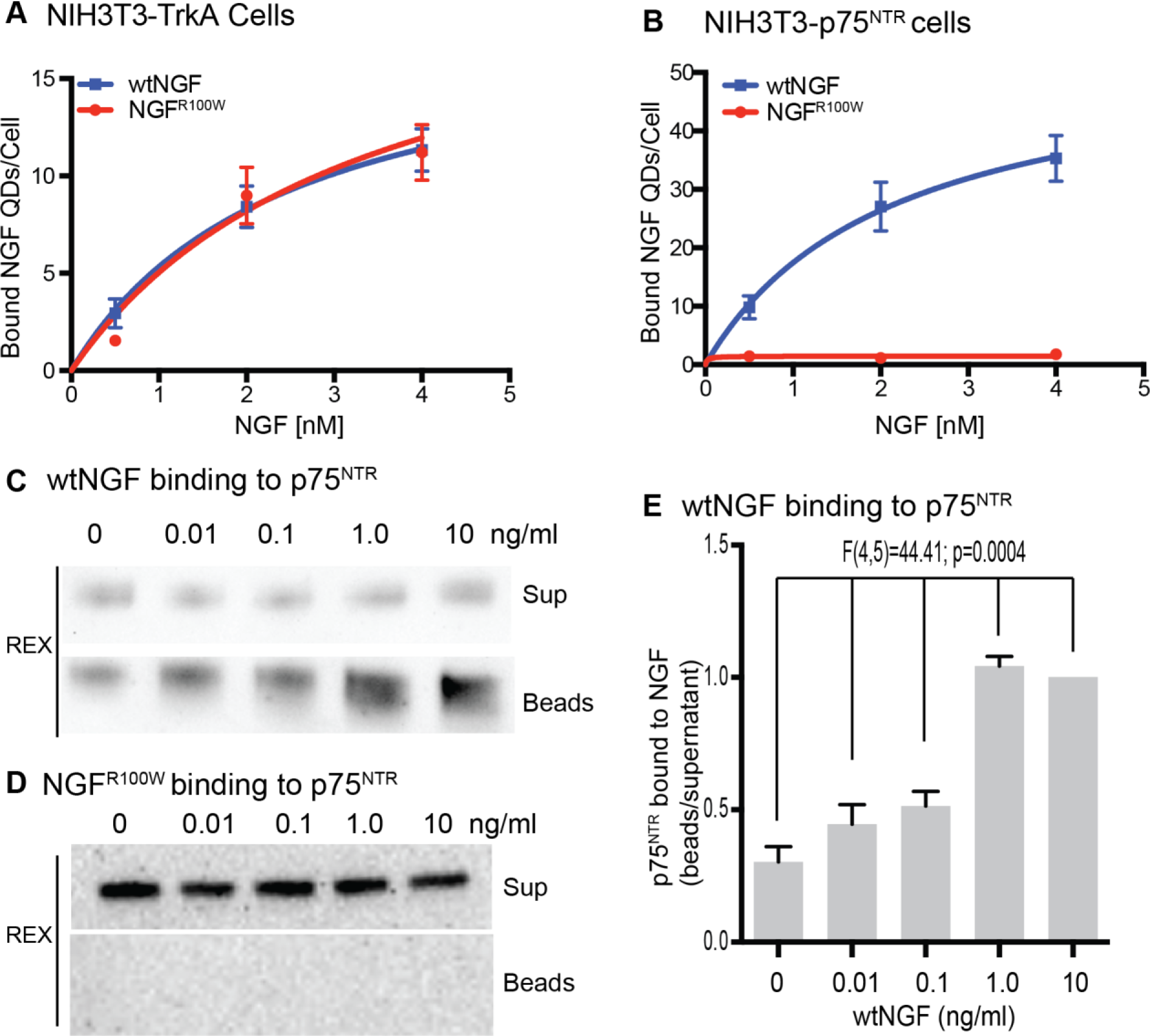
Classical hyperbolic saturation curves of wtNGF and NGF^R100W^ bound to TrkA or p75^NTR^-expressing NIH3T3 cells. Either NIH3T3 TrkA or p75^NTR^ cells were exposed to wtNGF-QD655/or NGF^R100W^-QD655 for 20-30 min at 20°C with range of different NGF-QD655 concentrations (**A, B**). Non-linear regression analysis using observed data was performed to examine the binding of NGF to the TrkA receptor (**A**) or p75^NTR^ (**B**). Blue squares and red circles represent surface-bound wtNGF and NGF^R100W^, respectively. Biotinylated wtNGF (**C**) or NGF^R100W^ (**D**) was incubated with recombinant extracellular domain of p75^NTR^ at 4°C for 2 hrs. The complex that contained biotinylated NGF was pulled-down using streptavidin-agarose (SA) beads at 4°C for 2 hrs. The beads were washed and boiled in SDS sample buffer. Both an aliquot of the supernatant (control) and the bead-bound samples were analyzed by Western blotting using a specific antibody against p75^NTR^(REX). Representative blot shows p75^NTR^ bound to wtNGF-SA beads (**C**) and p75^NTR^ bound to NGF^R100W^-SA beads (**D**). Relative REX signal from pulled-down beads was significantly increased with increased amount of wtNGF as shown in (**C**). ANOVA analysis was performed among different amount of wtNGF suggesting pulled down p75 ECD were significantly increased according to the increased amount of wtNGF (**E**) (F_(4,5)_=44.41, p=0.0004). In contrast, pulled down p75 ECD was not detectable even with 10 ngf of NGF^R100W^, only showed the level of baseline Rex signal intensity (**F**).

To further confirm that NGF^R100W^ binding to p75^NTR^ was markedly reduced, we carried out *in vitro* binding assays for p75^NTR^ using the extracellular domain (p75^NTR^-ECD). Either wtNGF or NGF^R100W^ in their biotinylated forms at a final concentration ranging from 0 to 0.25 nM was incubated with recombinant p75^NTR^-ECD; streptavidin-agarose beads were used to pull-down p75^NTR^-ECD that bound to biotinylated wtNGF or NGF^R100W^. The levels of p75^NTR^-ECD were assayed by immunoblotting using the anti-p75^NTR^ antibody, REX. **Fig 4C, E** shows that increasing concentrations of wtNGF resulted in increasing amounts of p75^NTR^-ECD in the pull-down complex. In contrast, NGF^R100W^ failed to pull down detectable p75-ECD even at the highest concentration (**Fig 4D**). We conclude that the binding affinity of NGF^R100W^ for p75^NTR^ is markedly reduced with respect to that for wtNGF; thus, NGF^R100W^ fails to bind p75^NTR^.

To extend these analyses, we then assayed if wtNGF and NGF^R100W^ differed with respect to TrkA-or p75^NTR^ receptor-mediated internalization. Since NGF^R100W^ showed reduced or absent binding to p75^NTR^, NGF^R100W^ would also fail to be internalized via p75^NTR^. We performed live cell imaging using NIH3T3-TrkA or NIH3T3-p75^NTR^ expressing cells (Huang et al., 1999). In addition to wtNGF and NGF^R100W^, we also took advantage of two well-characterized NGF mutants: 1) the KKE mutant which shows a significant decrease in binding affinity for p75^NTR^ but unaltered binding affinity for TrkA (Ibanez et al., 1992; Mahapatra et al., 2009); 2) the Δ9/13 mutant that poorly binds to TrkA while maintaining normal binding affinity for p75^NTR^ (Hughes et al., 2001). Accordingly, KKE and Δ 9/13 NGF served as positive controls for binding to TrkA and p75^NTR^, respectively. We produced monobiotinylated forms of KKE and Δ 9/13 NGF along with wtNGF and NGF^R100W^ to facilitate conjugation to QD655. These QD-655 labeled forms of NGF were incubated with cells at a final concentration of 0.2 nM at 37°C (30 min for NIH3T3-TrkA cells and 2 hrs for NIH3T3-p75^NTR^ cells) to allow for internalization. Incubations were followed by extensive washing at 4°C in PBS (three times). QD655 signals were captured by live imaging and defined as internal if the QD signal was found within the perimeter of the cell and at the same focal level as nucleus. As with wtNGF and the KKE mutant, NGF^R100W^ was internalized by NIH3T3-TrkA cells (**Fig 5A**). The process was receptor-mediated since premixing with 100x wtNGF (i.e. not conjugated to QD 655) eliminated NGF^R100W^-QD655 internalization. The Δ9/13 mutant were not internalized in NIH3T3-TrkA cells (**Fig 5A, C**). As predicted, NGF^R100W^ as well as the KKE mutant failed to be internalized into NIH3T3-p75^NTR^ cells; with both wtNGF and the Δ9/13 mutant, bright QD signals were detected inside NIH3T3-p75^NTR^ cells (**Fig 5B, C**). As a control, non-conjugated QDs were not detected in either cell lines. These results are further evidence that NGF^R100W^ is similar to wtNGF by binding to TrkA but differs in not binding to p75^NTR^.

**Figure 5.**
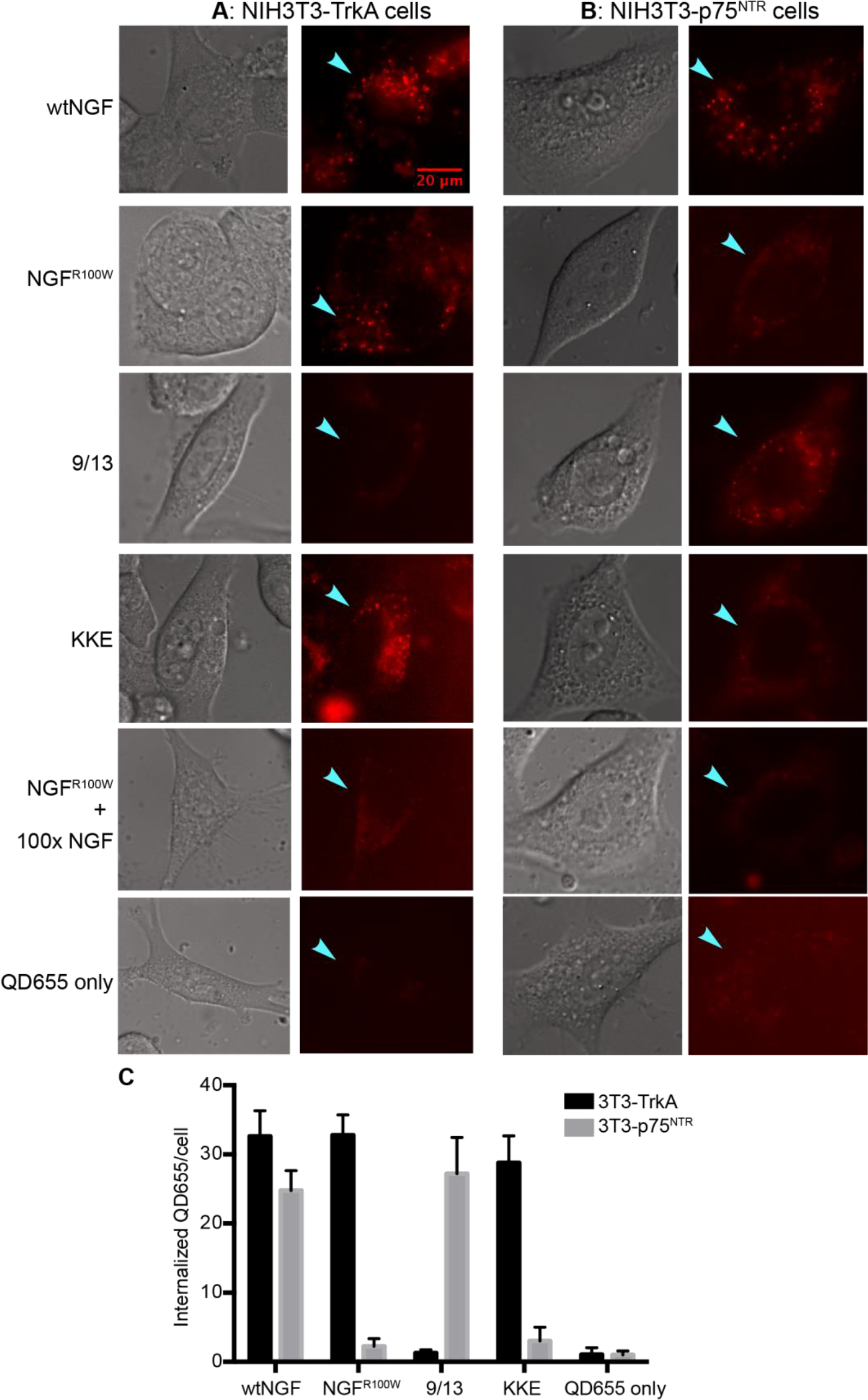
Imaging analysis of binding and internalization into NIH3T3-TrkA or –p75^NTR^ expressing cells. **A**: NIH3T3-TrkA cells were cultured on coverslips that were precoated with poly-L-lysine. Cells were rinsed and serum-starved for 2 hrs. Cells were then incubated with 0.2 nM of wtNGF-, or NGF^R100W^-, or KKE-or delta9/13-QD655-conjugates, or with QD655 alone for 30 min at 37°C. Following extensive rinses, QD655 signals were captured by live cell imaging. Representative images are shown. wtNGF, NGF^R100W^ and the KKE mutant that were used as positive controls for the TrkA receptor showed bright QD655 signals inside the cells. To ensure that internalization of NGF^R100W^ was receptor-mediated, we premixed NGF^R100W^ with 100 folds of unlabeled NGF prior to live imaging. The results show little internalization of NGF^R100W^, indicating the internalization of NGF^R100W^ into NIH3T3-TrkA cells was TrkA specific. **B**: Similarly, NIH3T3-p75^NTR^ cells were used to investigate internalization of the different forms of NGF proteins and representative images are shown after incubating with NGF-QD655 for 2 hrs at 37°C. The results show that both wtNGF and Δ9/13, that are known to bind to the p75^NTR^ receptor, were internalized into NIH3T3-p75^NTR^ cells and the QD655 signals were mostly concentrated around the peripheries of the cell. No signals were observed in the NIH3T3-p75^NTR^ cells when treated with either the KKE mutant or NGF^R100W^. **C**: internalized QD655 within the cells were quantitated.

### 5. NGF^R100W^ activates TrkA-mediated signaling pathways but failed to stimulate a p75^NTR^ downstream effector

Based on the binding and internalization results, we predicted that NGF^R100W^ activated TrkA-, but not p75^NTR^-mediated signaling pathways. We next determined if wtNGF and NGF^R100W^ activated two main effectors of TrkA downstream signaling cascades: Erk1/2 and Akt using PC12 cells (**Fig 6A**). For this purpose, PC12 was stimulated with 50ng/ml of either wtNGF or NGF^R100W^. Cells then were lysed and analyzed using immunoblotting. **Fig 6A, B, C** shows that NGF^R100W^ induced phosphorylation of TrkA, Erk1/2 and Akt to the extent comparable to wtNGF. We also performed semi-quantitative measurement of pTrkA activated by wtNGF or NGF^R100W^. The signals for pTrkA were normalized against GAPDH (Extended Fig 6-1). We did not detect a significant difference in pTrkA (Extended Fig 6-1). These findings suggest that NGF^R100W^ maintains its ability to activate TrkA, Erk 1/2, and Akt signaling pathways, which are primarily involved in providing trophic support for neuronal survival and differentiation.

**Figure 6.**
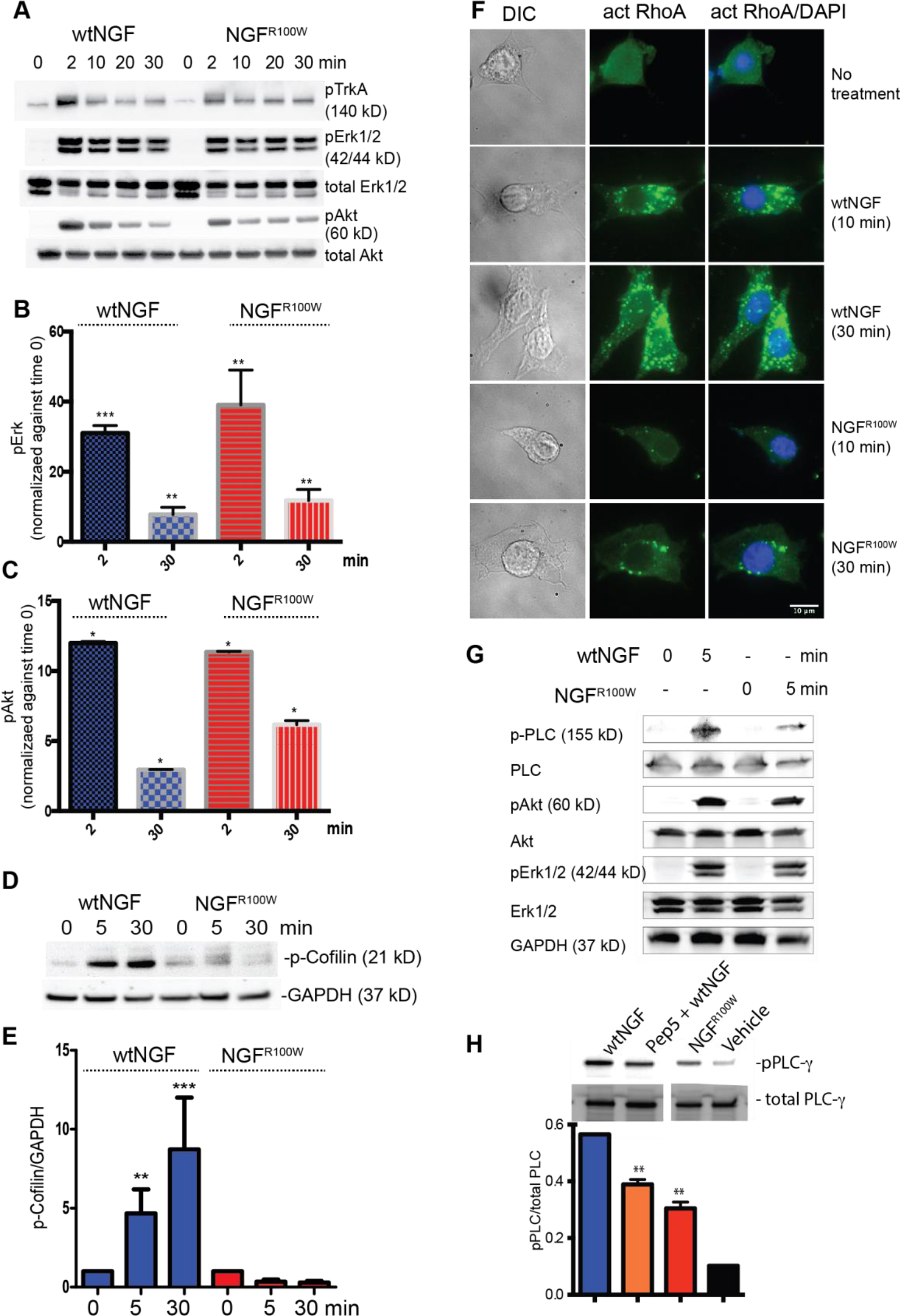
Analysis of TrkA and p75^NTR^-mediated signaling pathways. **A, B, C**: PC12 cells were serum-starved and were treated with 50ng/ml of either wtNGF or NGF^R100W^ for the indicated time intervals. Cell lysates were analyzed by SDS-PAGE and immunoblotting with specific antibodies as indicated. Treatment with either wtNGF or NGF^R100W^ induced phosphorylation i.e. activation of TrkA, Erk 1/2, and Akt. The blots were reprobed for total Akt or Erk1/2 as loading controls. We also measured the signals for pTrkA activated by wtNGF or NGF^R100W^. The signals for pTrkA were normalized against GAPDH (Extended Fig 6-1). **D, E**: Analysis of downstream signaling of p75^NTR^ in PC12 nnr5 by immunoblotting. The cells were prepared, treated and cell lysates analyzed by SDS-PAGE/immunoblotting with specific antibodies as in **A**. There was a significant reduction in phosphorylation of cofilin by NGF^R100W^ as compared to wtNGF. **F**: Immunostaining of RhoA-GTP in PC12 nnr5. PC12 nnr5 were plated on the coverslip coated by poly-L-lys, starved for 2 hr, treated by either wtNGF or NGF^R100W^, and the preparations were fixed, permeabilized followed by the protocol. DIC imaged cells, single staining for active form of RhoA (RhoA-GTP) or double staining for active RhoA and nucleus is shown. RhoA-GTP staining revealed RhoA activation was stronger in the cell treated with wtNGF than NGF^R100W^. **G, H:** Analysis of PLC-γ signaling in PC12 cells by immunoblotting. **G** showed PLC-γ stimulated by NGFR100W significantly differ from the one by wtNGF. In contrast, the same lysate showed similar amount of activation of Erk1/2 and Akt. PC12 cells were pretreated with the p75^NTR^ inhibitor Pep15 followed by treatment with 50 ng/ml NGF. In parallel samples, cells were treated with either vehicle, or 50 ng/ml with NGF or 50 ng/ml NGF^R100W^. Cell lysates were analyzed by SDS-PAGE/immunoblotting with specific antibodies as indicated. The data show thatactivation of PLC-γ was markedly suppressed by NGF^R100W^ compared to wtNGF. wtNGF failed to fully activate PLC-γ when p75^NTR^ is functionally inhibited, similar to partially activated PLC-γ when treated by NGFR100W.

We then assessed whether both wtNGF and NGF^R100W^ stimulated two signaling pathways downstream of p75^NTR^ receptors. We investigated if RhoA and Cofilin, two effectors downstream to p75^NTR^ (Yamashita and Tohyama, 2003; Vardouli et al., 2005), were phosphorylated or activated by wtNGF or NGF^R100W^. Using PC12^nnr5^ cells that express p75^NTR^ with little or no TrkA (Loeb and Greene, 1993), we tested if treatment of these cells with either wtNGF or NGF^R100W^ induced phosphorylation of Cofilin (i.e. p-Cofilin) by immunoblotting using an antibody that specifically recognizes the phosphorylated form of Cofilin. Our results show that Cofilin was activated when cells are treated by wtNGF. However, the level of phosphorylated Cofilin was significantly less in cells treated by NGF^R100W^ (**Fig 6D, E**). We then assayed for activation of RhoA, a signaling molecule upstream of Cofilin in the p75^NTR^ signaling cascades (Vardouli et al., 2005). As demonstrated by immunostaining with specific antibody to activated RhoA (i.e. RhoA^GTP^), RhoA^GTP^ exhibited marked activation by wtNGF as evident by strong cytosolic staining (**Fig 6F**). In contrast, NGF^R100W^ resulted in much less activation of RhoA^GTP^ with only sparse speckles of signals in the cytoplasm (**Fig 6F**). These data further confirm that NGF^R100W^ is ineffective in activating signaling cascades downstream of p75^NTR^.

Interestingly, we observed NGF^R100W^ was unable to fully induce phosphorylation of PLC-γ (**Fig 6G**). Therefore, we asked whether failure of activation of the p75^NTR^ signaling by NGF^R100W^ was responsible for the inability to fully phosphorylate PLC-γ. Indeed, there has been evidence suggesting that activated p75^NTR^ downstream effectors positively impact downstream of TrkA signaling (Ruiz-Arguello et al., 1996; Basanez et al., 1997; Ruiz-Arguello et al., 1998; Cremesti et al., 2002). For example, others have shown that ceramide-induced changes in membrane microenvironments facilitates PLC signaling. We therefore speculated that treating cells with wtNGF under conditions in which p75^NTR^ was inhibited would cause a reduction in PLC-γ activation, as shown in the case for NGF^R100W^ (**Fig 6H**). We used PC12 cells to examine if p75^NTR^ inhibition effected on PLC-γ by possible crosstalk between p75^NTR^ and TrkA. Our results showed that phosphorylation of PLC-γ was decreased when PC12 were pre-treated with a p75^NTR^ inhibitor, TAT-pep5(Head et al., 2009), followed by stimulation with wtNGF (Hasegawa et al., 2004), as shown in **Fig 6F**. These results suggest that failure of p75^NTR^ signaling by NGF^R100W^ leads to failure of PLC-γ signaling by as yet unknown mechanisms, even though NGF^R100W^ stimulates other downstream signaling such as Erk1/2 and Akt under TrkA. Taken together, NGF^R100W^ activates most TrkA downstream signaling events but not those mediated by p75.

### 6. NGF^R100W^ was retrogradely transported in a fashion similar to wtNGF

Axons feature prominently in facilitating trafficking and signaling of NGF. A possibility existed that the loss of pain signaling in NGF^R100W^ was due to its inability to be effectively transported retrogradely to the cell soma in sensory neurons. The ability of R100W to induce priming suggested that it was effectively transported retrograde in axons to engage the cell body responses that support priming. To test this possibility, we established microfluidic cultures of dissociated rat embryonic E15.5 DRG neurons. To visualize trafficking of NGF, we conjugated biotinylated wtNGF or NGF^R100W^ with streptavidin-QD605 at 1 NGF dimer: 1 QD605 ratio. This experimental paradigm was used to track axonal movement of a single NGF dimer by live imaging to produce highly quantitative results (Sung et al., 2011). wtNGF was retrogradely transported with an average speed of 1.5 μm/sec (**Fig 7A**), which is consistent with previous results (Cui et al., 2007; Sung et al., 2011). Based on the analysis of more than 100 endosomes/condition, the average moving speed of NGF^R100W^ revealed no marked difference from that of wtNGF (**Fig 7B**).

**Figure 7.**
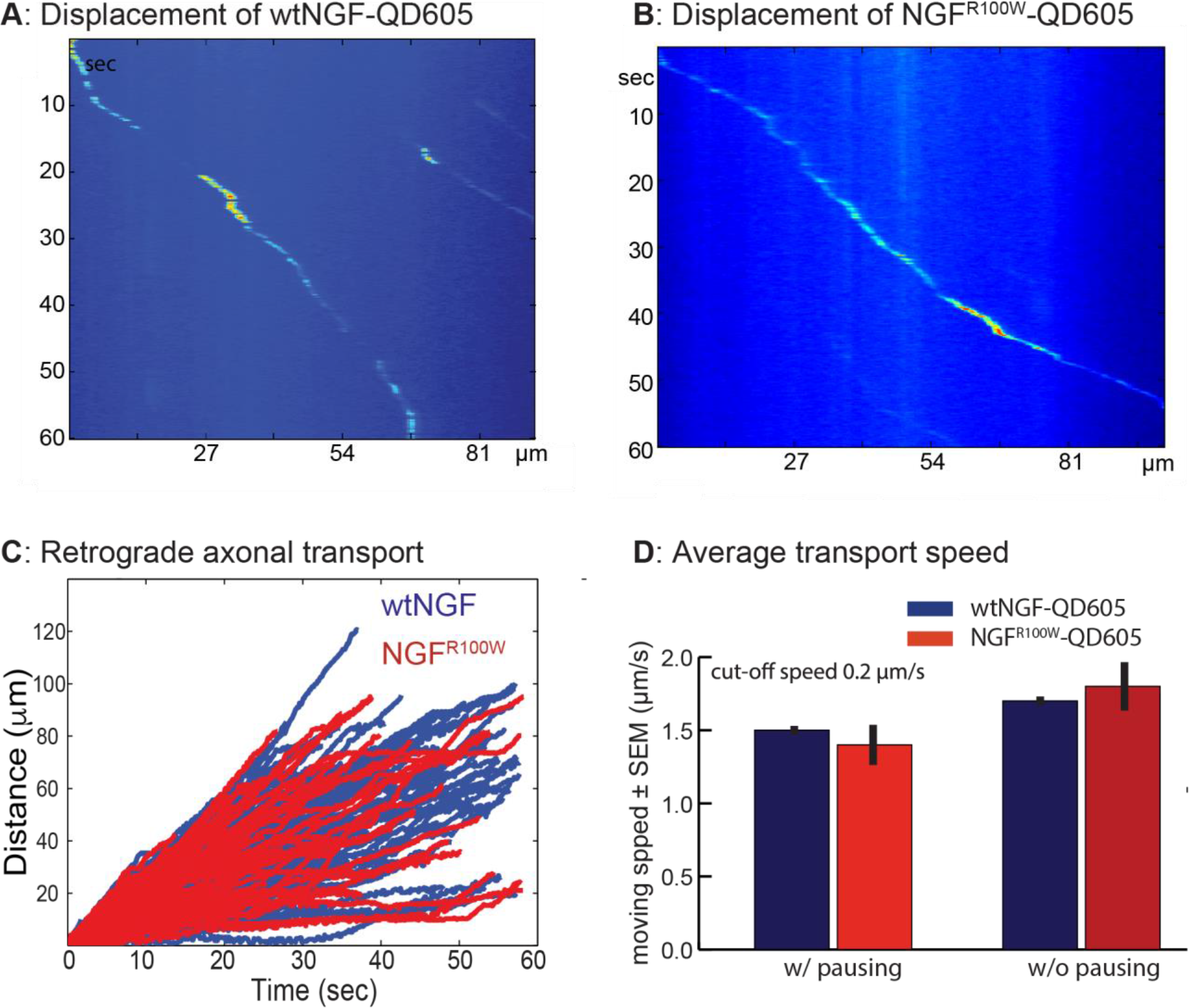
Analysis of retrograde axonal transport by live imaging. **A, B**: rat E15.5 DRG neurons were cultured in microfluidic chamber. Kymographs of axonal movement of wtNGF and NGF^R100W^ based on real-time imaging series of axonal transport assays. The graphs represent spatial position of QD605 signals (in μm) over time (seconds). The results for both wtNGF and NGF^R100W^ showed similar slopes, suggesting NGF^R100W^ moves at a speed within the axon similar to that of wtNGF. (**C**) Overlayed kymographs of displacement of axonal QD605 signals for wtNGF and NGF^R100W^. 100-150 QD605 signals for either wtNGF (blue) or NGF^R100W^ (red) were analyzed and superimposed. The results demonstrate that axonal movement of wtNGF and NGF^R100W^ behaves in a strikingly similar fashion. (**D**) The total average transport speeds including pausing for wtNGF and NGF^R100W^ were calculated to be 1.5 mm/s and 1.4 mm/s, respectively. The moving velocity without pausing i.e. during the “go” motion period was 1.7 mm/s for both wtNGF and NGF^R100W^.

NGF is known to exhibit a ‘go and stop’ behavior during transport (Cui et al., 2007; Sung et al., 2011). Therefore, we tested if the moving speed during the ‘go’ period differed significantly between wtNGF and NGF^R100W^. Our results demonstrated that this was not the case (**Fig 7D**). By superimposing the kymographs of NGF^R100W^ onto those wtNGF (**Fig 7C**), our results confirmed that NGF^R100W^ behaved exactly like wtNGF during retrograde transit from the axonal terminal to the cell body. We thus conclude that the critical function in retrograde axonal transport of NGF in sensory neurons is preserved in NGF^R100W^.

### 7. Study of contribution and TrkA or p75^NTR^ to NGF-induced sensitization effect in vivo

Our findings suggest that NGF^R100W^ retained its ability to bind to and activate TrkA while failing to engage p75^NTR^. Yet when injected into adult rats, NGF^R100W^ still induced hyperalgesic priming without causing acute sensitization ot mechanical stimuli. These results raise the possibility that TrkA and p75^NTR^ may play distinct role(s) in NGF-induced hyperalgesia and hyperalgesic priming. To this end, we used pharmacological reagents to selectively inhibit signaling downstream of TrkA and/or p75^NTR^: K252a for blocking TrkA activation and GW4869 for inhibiting neutral sphingomyelinase downstream of p75^NTR^.

Vehicles or inhibitors (K252a, GW4869, K252a + GW4869) were injected only once at 5 min prior to NGF injection (**Fig 8A**). We then measured the mechanical nociceptive threshold at 1 hr and 7 days after NGF injection as outlined in the experimental design (**Fig 8A**). K252a produced a robust reduction (~30%) in NGF hyperalgesia (p=0.0001); GW4869 had a small but insignificant reduction (~7%, p=0.0557); combination of K252a with GW4869 caused a ~25% reduction (p<0.0001). Consistent with results in presented Fig 1, NGF-induced hyperalgesia had returned to baseline nociceptive threshold at Day 7 following NGF treatment, regardless of inhibitors (**Fig 8A**).

**Figure 8.**
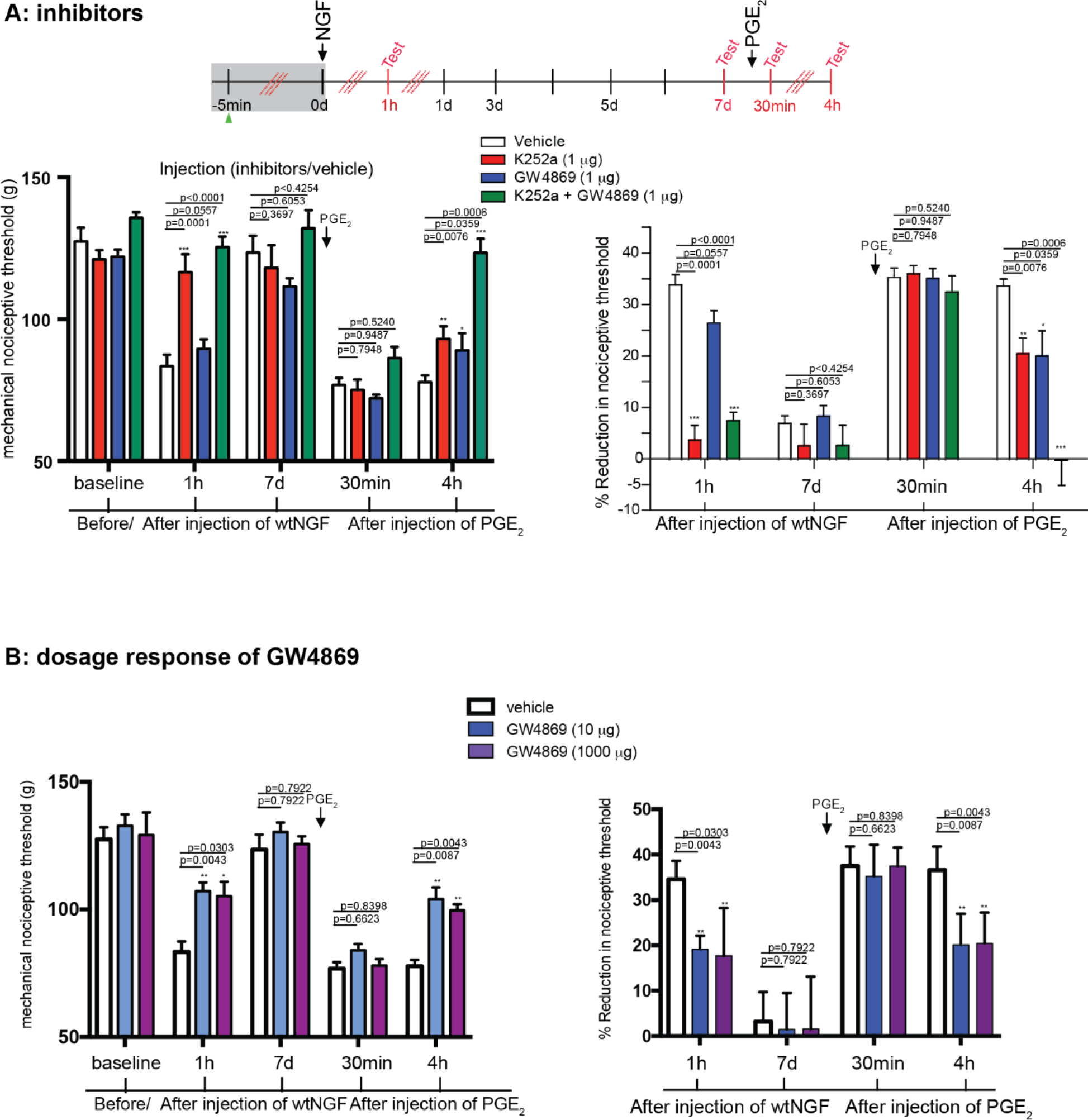
Role of TrkA and p75^NTR^ in nociceptive response and hyperalgesic priming. **A**: 5 μl (1 μg) of K252a, GW4869 or vehicle (10% DMSO) were administered intradermally on the dorsum of the hind paw of adult rats at the same site where NGF was going to be injected. Flowing NGF injection (200 ng of wtNGF), the Randall-Selitto method was used to measure mechanical hyperalgesia as in A. n=4 for vehicle, K252a, GW4869; n=6 for K252a + GW4869. Data are presented as mean + SEM. p values were calculated using unpaired t-test. *** indicates when p<0.001. **B:** Higher dosages of GW4869, 10μg (n=4) and 1000μg (n=4), were injected intradermally on the dorsum of the hind paw of adult rats following by NGF injection as in **A**. Same method was applied to measure mechanical hyperalgesia used as in **A**. Data are presented as mean + SEM. p values were calculated using unpaired t-test.

We then measured the impact of these inhibitor treatment on priming following injection of PEG_2_ as in Figure 1. Treatment with K252a, GW4869 or both did not significantly change acute sensitization 30 min following PEG_2_ injection (**Fig 8A**). However, at 4 hrs following PEG_2_ injection, K252a and GW4869 each alone produced a moderate reduction PGE_2_ hyperalgesia (p=0.0076 for K252a; p0.0359 for GW4869); Strikingly, combination of K252a with GW4869 induced complete elimination of PGE_2_ hyperalgesia (p=0.0006) (**Fig 8A**). These results further support an interaction between TrkA and p75^NTR^ in mediating the priming effect by NGF.

Since there is discrepancy between NGF^R100W^ injection studies (**Fig 1B,C**) versus GW4869 treatment in inducing acute hypersensitivity (**Fig 8A**), we repeated Randall-Selitto experiment with higher dosages of GW 4869. We chose two different dosages, 10μg and 1000μg of GW 4869. Previous studies have demonstrated that 11.55μg/injection of GW 4869 was effective in blocking acute sensitization by NGF(Khodorova et al., 2013, 2017). The 1000μg dose was to ensure a maximal effect was achieved. As demonstrated in **Fig 8B**, 10 μg/injection appeared to yield a maximal effect; both dosage, 10 μg and 1000 μg successfully prevented NGF-induced acute hyperalgesia at 1hr after NGF injection, which is in line with published studies(Khodorova et al., 2013, 2017) but also from our own studies presented in **Fig 1B,C**. At 1hr after NGF injection, while vehicle injection decreased the mechanical pain threshold about 34.57%, 10μg and 1000μg of GW4869 decreased the threshold about 19.14% (p=0.0043) and 17.64% (p=0.0303) respectively. (**Fig 8B**). We then measured the priming following injection of PGE_2_ with these two high dosages of GW4869. Both dosages also blocked priming significantly compared to vehicle treated (p=0.0087 and p=0.0043) (**Fig 8B**). These data further confirmed our result pointing to the loss of p75^NTR^ downstream signaling in NGF^R100W^ as a mechanism underlying loss of pain perception. Taken together, these data point to both TrkA and p75^NTR^ responsible in mediating acute hypersensitivity and priming.

## Discussion

Studies on a naturally occurring mutation in HSAN V patients, NGF^R100W^, have provided new insights into NGF-mediation of nociceptor function. Using molecular, cellular, and biochemical techniques we demonstrated that NGF^R100W^ retained the ability to bind and signal through TrkA receptors and robustly support the trophic status of DRGs. In contrast, NGF^R100W^ failed to bind or activate of p75^NTR^. Our *in vivo* studies have demonstrated that inhibition of TrkA and p75^NTR^ signaling, as well as delivery of antisense reagents targeting these receptors, provided evidence for a role for both receptors in nociceptor regulation. While TrkA activation mediated acute sensitization and priming, p75^NTR^ signaling appeared to contribute only to priming, thereby augmenting the TrkA response. Remarkably, and in contrast to its ability to induce trophic responses *via* TrkA, NGF^R100W^ induced nociceptor priming but differed from wtNGF by not causing acute sensitization. These observations are evidence for distinct TrkA signaling pathways mediating trophic effects and nociceptor function and provide insights into the biology of HSAN V.

Consistent with previous results (Covaceuszach et al., 2010; Capsoni et al., 2011), we confirmed that while NGF^R100W^ binding and activation of TrkA is robust, binding and signaling through p75^NTR^ is essentially absent. Important to its ability to prime nociceptors, NGF^R100W^ was internalized at axonal tips of DRG sensory neurons and traveled toward the cell body at the speed and with the velocity characteristic of wtNGF. Given properties in common with wtNGF, it is unclear why NGF^R100W^ failed to induce acute sensitization. Possibilities include differences with respect to wtNGF in the structure of the NGF^R100W^/TrkA complex, the structure of the complex in signaling endosomes, or the downstream partners recruited to these complexes. Furthermore, it is hard to exclude the possibility that TrkA dowstrenam that is implicated in pain could be negatively affected by intertwined crosstalk between p75^NTR^ and TrkA as shown in partial PLC gamma activation by NGF^R100W^. It suggests possibility that TrkA downstream that supports trophic function is retained in NGF^R100W^ mutation while p75 downstream and partial TrkA downstream that affects pain threshold is knocked-downed in NGF^R100W^. In any case, it is apparent that differences must exist for the signaling events that subserve TrkA trophic and its acute effects on nociceptor sensitization. Further studies to decipher the basis for these differences will benefit from the ability to compare the signaling properties of NGF^R100W^/TrkA with wtNGF/TrkA.

Our data helps to explain the clinical manifestations of NGF^R100W^ mutation. Studies of affected families point to considerable inter-patient variability. Both homozygous and heterozygous individuals demonstrate orthopedic manifestations, the most frequent of which is multiple painless fractures, typically in the legs and feet(Einarsdottir et al., 2004; Minde et al., 2004; Larsson et al., 2009; Capsoni, 2014). Homozygous patients show decreased pain sensation, mainly at the forearms and legs. In contrast, they respond to truncal pain and register visceral pain. Distal testing of temperature thresholds showed increases in some homozygous and heterozygous patients. Sensitivity to soft touch, joint position, vibration sensation and visceral pain are normal. Sural nerve biopsy reveals loss of C and A delta sensory fibers in both homozygotes and heterozygotes(Minde et al., 2004; Minde, 2006; Minde et al., 2009; Sagafos et al., 2016). All patients show reduced sensory innervation of skin and reduced sympathetic innervation of sweat glands, with more marked changes in homozygotes ((Axelsson et al., 2009). While a decrease in pain is consistent with the inability of NGF^R100W^ to induce acute nociceptor sensitization, the clinical picture is that of a length-dependent sensory and sympathetic neuropathy. Indeed, this explains best the painless fractures and decreased pain sensation in distal lower limbs, increased distal thresholds for cold and heat perception, loss of sensory and sympathetic innervation of skin and preservation of truncal pain. In view of the preservation of TrkA signaling by NGF^R100W^, the question arises as to how NGF^R100W^ causes the syndrome. The likely cause is failure to secrete the protein in sufficient amounts to support the distal axons of sensory and sympathetic neurons. NGF is critical for the survival and maintenance of sympathetic and sensory neurons (Ibanez et al., 1992; Kew et al., 1996; Casaccia-Bonnefil et al., 1998; Sofroniew et al., 2001; Chao, 2003). Decreased secretion of NGF^R100W^ as demonstrated in other studies must be confirmed with human cells expressing the mutant protein (Larsson et al., 2009; Covaceuszach et al., 2010).

A causal link between NGF deficiency in innervated targets and neuronal dysfunction and degeneration was established during the earliest studies on NGF (Levi-Montalcini and Hamburger, 1951; Cohen and Levi-Montalcini, 1956; Levi-Montalcini and Angeletti, 1961, 1963; Levi-Montalcini, 1964) and has been shown in studies of the developing and mature nervous system. Attempts to increase NGF availability in hopes of reducing neurodegeneration (Wilcox and Johnson, 1988; Apfel et al., 1994; Apfel and Kessler, 1995, 1996; Apfel, 1999a, b; McArthur et al., 2000; Apfel, 2002; Cattaneo et al., 2008) have failed in part due to NGF dose-limiting pain (Eriksdotter Jonhagen et al., 1998; Apfel, 2000, 2002). That NGF^R100W^ maintains trophic functions without acutely sensitizing nociceptors has suggested that this isoform of NGF might be used to provide trophic support without causing pain (Capsoni et al., 2011; Capsoni, 2014). That NGF^R100W^ maintains the ability of wt NGF to prime nociceptors raises the caution that NGF^R100W^ treatment may not avoid fully the pain induced by wild type NGF.

Increasing evidence supports that both TrkA-and p75^NTR^-mediated signaling pathways are intimately involved in NGF-induced hyperalgesia (Nicol and Vasko, 2007). Extensive human genetic studies strongly support an essential role played by TrkA in pain sensation; mutations in TrkA that result in loss or reduced TrkA activity are associated with congenital insensitivity to pain with anhidrosis (CIPA) (Indo, 2001, 2002). Additionally, inhibiting of TrkA-mediated signaling pathways such as Erk1/2 (Aley et al., 2001; Dai et al., 2002), the PI3K/Akt (Zhuang et al., 2004) and the PLC-γ (Chuang et al., 2001) has been shown to block NGF-induced sensitization, both *in vivo* and *in vitro*. Consistent with these findings, we used specific antisense oligos to attenuate expression of TrkA *via* intrathecal administration or administered K252a to inhibit TrkA activation via acute intraplantar injection. Both approaches affirmed that TrkA is required for both the acute and chronic phase of sensitization induced by NGF.

Unlike TrkA, a role for p75^NTR^ in NGF-induced hyperalgesia has been implicated largely by indirect evidence: 1) intrathecal administration of anti-p75^NTR^ into animals reduced temperature hyperalgesia and mechanical allodynia after nerve injury (Obata et al., 2006); 2) direct application of an p75^NTR^ antibody to a crushed sciatic nerve suppressed mechanical allodynia (Fukui et al., 2010); 3) pretreatment with a p75^NTR^ antibody prevented the increase in the number of action potentials induced by NGF (Zhang and Nicol, 2004); 4) intraplantar injection of proNGF, which selectively activates p75^NTR^, and not TrkA, induced hyperalgesia(Khodorova et al., 2013); 5) NGF-induced sensitization was attenuated by inhibiting p75^NTR^-mediated activation of the sphingomyelin-ceramide-sphingosine 1 phosphate and the c-JUN kinase pathway (Zhang et al., 2002; Doya et al., 2005; Obata et al., 2006; Zhang et al., 2006; Khodorova et al., 2013).

We confirmed a role for p75^NTR^ in acute sensitization of mechanical nociceptors by using GW4869. However, a discrepancy exists between priming studies of NGF^R100W^ in Fig 1 versus GW4869 in Fig 8A; NGF^R100W^ induced a priming effect comparably to wtNGF, while GW4869 failed to block the priming effect by wtNGF even at higher dosages (Fig 8B). This could be explained by the lack of specificity of GW4869 in blocking neutral SMase2(Canals et al., 2011). GW4869 specificity issue was also raised in the literatures showing that GW4869 not only block SMase but also PLC, PP2A(Luberto et al., 2002). Moreover, previous literature has showed that GW4869 abrogated NGF mediated TrkA trophic effects on cell viability(Candalija et al., 2014) suggesting block of nSMase2 could lead partial block of TrkA mediated trophic signaling such as pAkt (Gills et al., 2012). Given all these, GW4869 effect could not be the same as loss of p75 downstream by NGF^R100W^. This may explain the inconsistency in our data showing incapability of priming by GW4869 versus the priming capability by NGF^R100W^.

We are also aware of the specificity issues associated with the use of K252a inhibitor to block TrkA. K252a selectivity has been studied in many literatures. For example, K252A is known to inhibit PKCs, which is also implicated in p75 downstream (Mizuno et al., 1993). Not only inhibiting TrkA, at certain range of concentration, K252a was also found to act as partial inhibitor of the PDGF receptor (Nye et al., 1992).

Given all the caveats associated with the use of inhibitors in our studies, our data suggest that p75^NTR^ plays an important role in NGF-induced pain function(Zhang and Nicol, 2004; Watanabe et al., 2008; Iwakura et al., 2010; Khodorova et al., 2013, 2017). When p75 inhibition is combined with TrkA suppression, both acute response and priming resulted in a greater reduction in NGF-mediated hyperalgesic priming. These results support a role for p75^NTR^ that is most evident in its ability to synergize with TrkA to mediate NGF-induced hyperalgesic priming. Therefore, the nociceptive functions of NGF include contributions from both TrkA and/or p75^NTR^.

To avoid the pitfalls associated with the use of inhibitors for TrkA or p75NTR-signaling pathways, future studies using novel genetic models are needed to further dissect the contributions of TrkA and p75^NTR^ to NGF-induced sensitization. For example, TrkA activity can be specifically inhibited by injection of nanomolar concentrations of derivatives of the general kinase inhibitor PP1(1NMPP1 or 1NaPP1) to block NGF signaling in TrkA(F592A)-knock in mice (Chen et al., 2005). Ablation of p75^NTR^ can be induced in a conditional p75^NTR^ knockout mice (Bogenmann et al., 2011; Wehner et al., 2016). Use of these models is expected to further clarify the contributions of TrkA and p75^NTR^, and the possibility that their signaling pathways interact to effect NGF-induced sensitization.

## Acknowledgement

We would like to thank Pauline Hu for assistance in DRG cultures, Dr. Chihye Chung for assisting with the single cell patch clamp experiments, Ms. R. Shibata and N. D. Storslett for their assistance in protein purification. Funding supports are from NIH (PN2 EY016525), Down Syndrome Research and Treatment Foundation, Larry L. Hillblom Foundation, Alzheimer’s Association, Thrasher Research Fund (WCM); NIH UCSD ADRC P50 Pilot grant, Larry L. Hillblom Foundation startup grant, Tauopathy Foundation (CW) and R01 NS084545 (JDL); NIH UCSD T32 Neuroplasticity of Aging Training Grant (KS).

## Author Contributions

KS, LFF, WY, XZ, KZ, MLP, MTM and CW performed the experiments; KN contributed to KKE and 9/13 constructs; KS, BC, WCM, JDL and CW wrote the manuscript.

---

**Extended Figure Legend (Figure 6-1).**
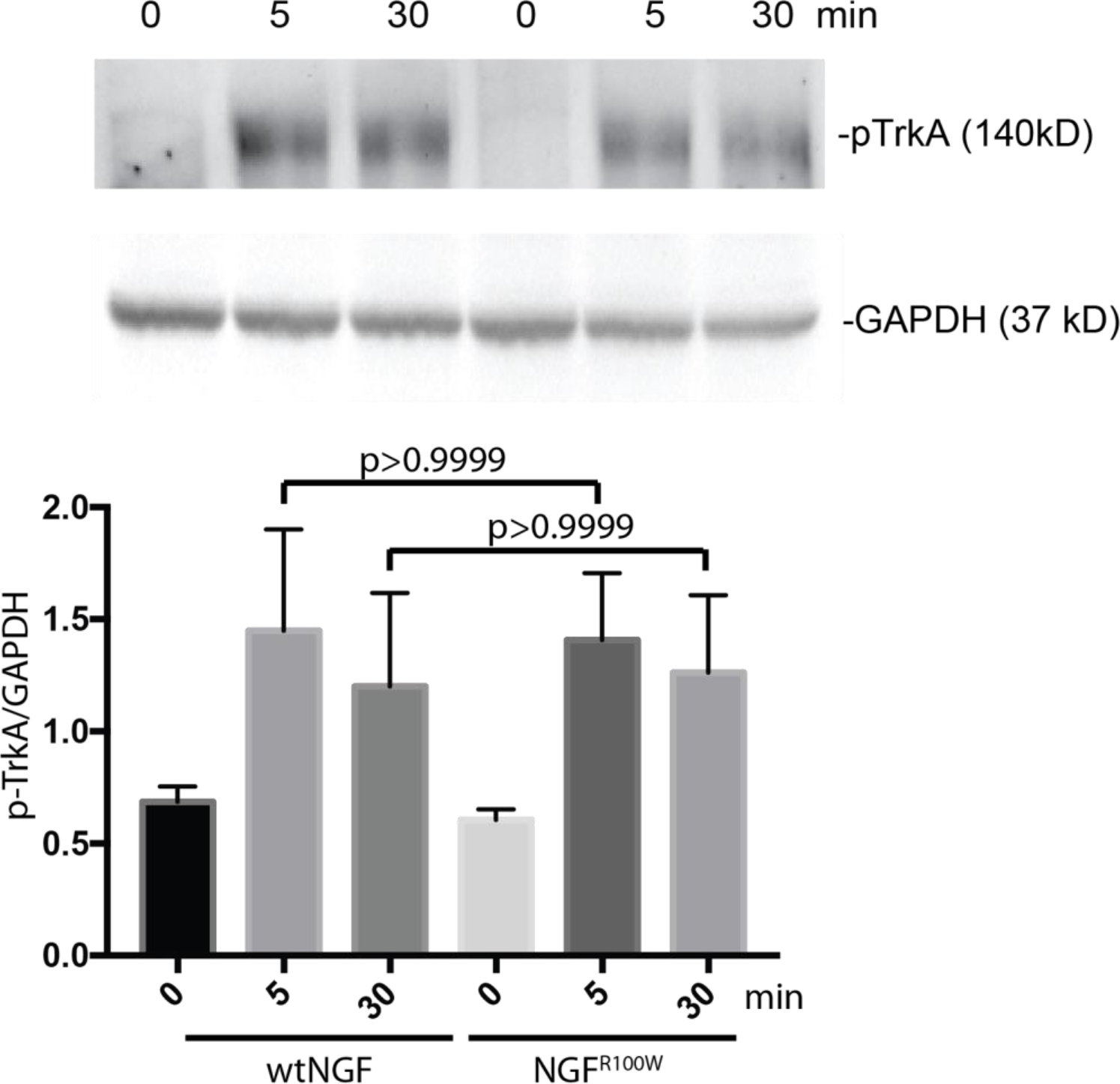
PC12 cells were serum-starved and were treated with 50ng/ml of either wtNGF or NGF^R100W^ for the indicated time intervals. Cell lysates were immunoprecipitated with a rabbit anti-TrkA antibody (UBI: Cat# 06-574, 1μg) and the immunoprecipitates were analyzed by SDS-PAGE/immunoblotting with a mouse anti-phopsho-Tyrosine antibody (Cell Signaling, Cat#9411, 1/200). An aliquot from each lysate was also analyzed by SDS-PAGE/immunoblotting with a mouse anti-GAPDH antibody (GenTex, GTX627408). The experiments were repeated for at least three times independently. The signals for pTrkA signals (140 kD) were quantitated using Bio-Rab ImageLab and were normalized against GAPDH (37 kD). Representative images and quantitative data are presented.

